# Unique soil fungal communities are associated with disappearing ash trees in a northern temperate hardwood forest

**DOI:** 10.64898/2026.07.11.737980

**Authors:** Farrar R. Ransom, Paul Metzler, Elizabeth Studer, Matthew P. Ayres, V. Bala Chaudhary

## Abstract

Native ash trees are destined for functional extinction in North America due to the spread of the non-native emerald ash borer. Yet, the consequences of ash loss for soil fungi are unclear. To address this, we employed a factorial study of forest soil fungi in two hydropedological soil types beneath four canopy tree species -- including white ash (*Fraxinus americana*). Sporocarp surveys and community DNA metabarcoding from soil samples revealed patterns in fungal communities related to canopy tree species but not soil type. Ash trees supported a particularly rich soil fungal community that was distinguishable from communities beneath beech, birch, and maple. We identified over 100 fungal taxa (OTUs) that are at risk of decline or loss from the studied forest, due to their association with ash. Our results indicate that canopy tree species influence soil fungi much more broadly than just the species with which they have mycorrhizal associations.

## Introduction

Rapid increases in the number of extinctions over the past century places species loss among the greatest ecological concerns of our time (Barnosky et al., 2011; IPBES, 2019). Coextinctions of additional species following the primary loss of a partner species complicate extinction impacts (Colwell et al., 2012). Although there are now over 400 fungal species listed as vulnerable or endangered on The IUCN Red List of Threatened Species (2025), the cryptic nature of fungi impedes extinction risk assessments (Dahlberg and Mueller, 2011; van der Linde et al., 2012; Nic Lughadha et al., 2020; Mueller et al., 2022). As many fungi are obligate symbionts of plants, risk of fungal coextinction is high (May et al., 2019; Niskanen et al., 2023). While coextinction risk could be utilized to sidestep challenges in quantifying population size when evaluating fungal vulnerability (Moir and Brennan, 2020; Mueller et al., 2022), host identity and specificity are still unclear for many fungi (Nic Lughadha et al., 2020; Niskanen et al., 2023). Prospects of fungal extinction increase the importance of documenting fungal species and their ecology, especially their host associations.

Invasive species are a notable driver of plant and animal extinctions, contributing to 25% of all plant extinctions in recent centuries (Blackburn et al., 2019; Nic Lughadha et al., 2020). The emerald ash borer is among the most damaging invasive insects yet for North American forests (Herms and McCullough, 2014). Since the introduction of the emerald ash borer in the late 1990s, ash trees have rapidly reached a state of functional extinction in many regions across the US and Canada (Klooster et al., 2018). In Michigan, near the epicenter of the invasion, ash mortality reached over 99% by 2009, reducing populations to the point that ash trees no longer contribute significant ecosystem services (Flower et al., 2013; Klooster et al., 2014). As the tree composition of these forests shift, the abundance of other organisms, including fungi, are expected to shift as well (Bellard et al., 2012; Klooster et al., 2018). Fungi are essential to forest ecosystems, especially in their contributions to plant health (van der Heijden et al., 2015; Anthony et al., 2022) and nutrient cycling (van der Wal et al., 2013; Fernandez and Kennedy, 2016). A better understanding of how fungal communities interact with ash and other tree species is necessary for projecting how northern forests will change in the coming decades.

Plant communities unequivocally shape fungal communities, but the importance of tree species identity relative to other variables remains unclear (Talbot et al., 2014; Hicks Pries et al., 2023). Tree species are often assessed by their dominant mycorrhizal fungi association, most commonly arbuscular mycorrhizal (AM) and ectomycorrhizal (EcM). AM and EcM fungi tend to be associated with different clades of plant species, with EcM fungi usually specializing to a greater extent (Toju et al., 2014; Zhu et al., 2018). Tree mycorrhizal association influences fungal community structure and function (Fernandez and Kennedy, 2016; Lin et al., 2022), but recent studies have begun to uncover differences in fungal communities among trees with the same mycorrhizal type (van der Linde et al., 2018; Hicks Pries et al., 2023; Fitch et al., 2023). These studies employ various methods to control for compounding effects of environmental variables such as is hydropedological soil type, which can also shape fungal communities (Talbot et al., 2014; Tedersoo et al., 2016; Steidinger et al., 2020). While Fitch et al. (2023) found distinct fungal communities among tree species, including AM associated white ash and white cedar, ash trees were not compared to another AM associated deciduous tree species. Further investigation beyond general tree mycorrhizal association is required for discerning how the loss of ash may impact fungal communities.

In advance of the impending loss of ash trees, we compared soil fungal communities with a factorial study in a northern-temperate forest, examining sixty replicate plots with different canopy tree species and soil types. To characterize the fungal community, we analyzed fungal community DNA extracted from soil samples. Field surveys of fungal sporocarps (reproductive fruiting bodies) were also conducted, as fungal metabarcoding analysis cannot distinguish between active fungi, dormant fungal spores, and dead fungi (van der Linde et al., 2018). Our investigations considered four tree species: white ash (*Fraxinus americana*), sugar maple (*Acer saccharum*), American beech (*Fagus grandifolia*), and yellow birch (*Betula alleghaniensis*). White ash and sugar maple associate with AM fungi while yellow birch and American beech associate with EcM fungi (Zhu et al., 2018). Plots of these trees were established across two common podzol soil types. Therefore, possible results of our study included patterns in forest fungal communities due to tree species, soil type, and any cross interaction (tree species x soil type). At minimum, we expected community differences to arise among tree species according to mycorrhizal association type. The factorial design, sequencing choices, and sporocarp surveys included in this study make it one of the most robust investigations to date on soil fungal communities under different tree species.

## Materials and Methods

### Study Site

Our study was conducted in the Hubbard Brook Experimental Forest (HBEF) within the White Mountain National Forest in central New Hampshire. HBEF consists of a 3,179-hectare valley bordered by two mountains and ridge lines to the north and south (Holmes and Likens, 2016a). The White Mountain region and HBEF are within the unceded homelands of the Indigenous Abenaki peoples. The forest was last logged in the late 1800s to early 1900s and, at the time of our study, was regarded as a mature second-growth northern hardwood forest (Siccama et al., 2007; Battles et al., 2014). Typical to the region, a northern hardwood forest ecosystem covers the lower slopes transitioning to spruce-fir forest at higher elevations. The canopy trees in our study area were chiefly American beech, sugar maple, and yellow birch, with occasional groves that included white ash (van Doorn et al., 2011). The emerald ash borer was discovered within HBEF in 2021 and had not yet begun affecting ash trees at the time of our study.

Data were collected from 60 plots previously established by Studer (2023). Each plot was circular with a 10-meter radius and flags marking the center and the four cardinal directions along the edge (Figure 1). Each plot was dominated by one of four tree species: white ash, sugar maple, American beech, or yellow birch (n = 15 for each). In all plots, the dominant tree species contributed > 50% of the total basal area and the other three study species each made up < 10% of the basal area. Stems with less than 10 cm2 basal area were not included in the plot basal area calculations.

**Figure 1.**
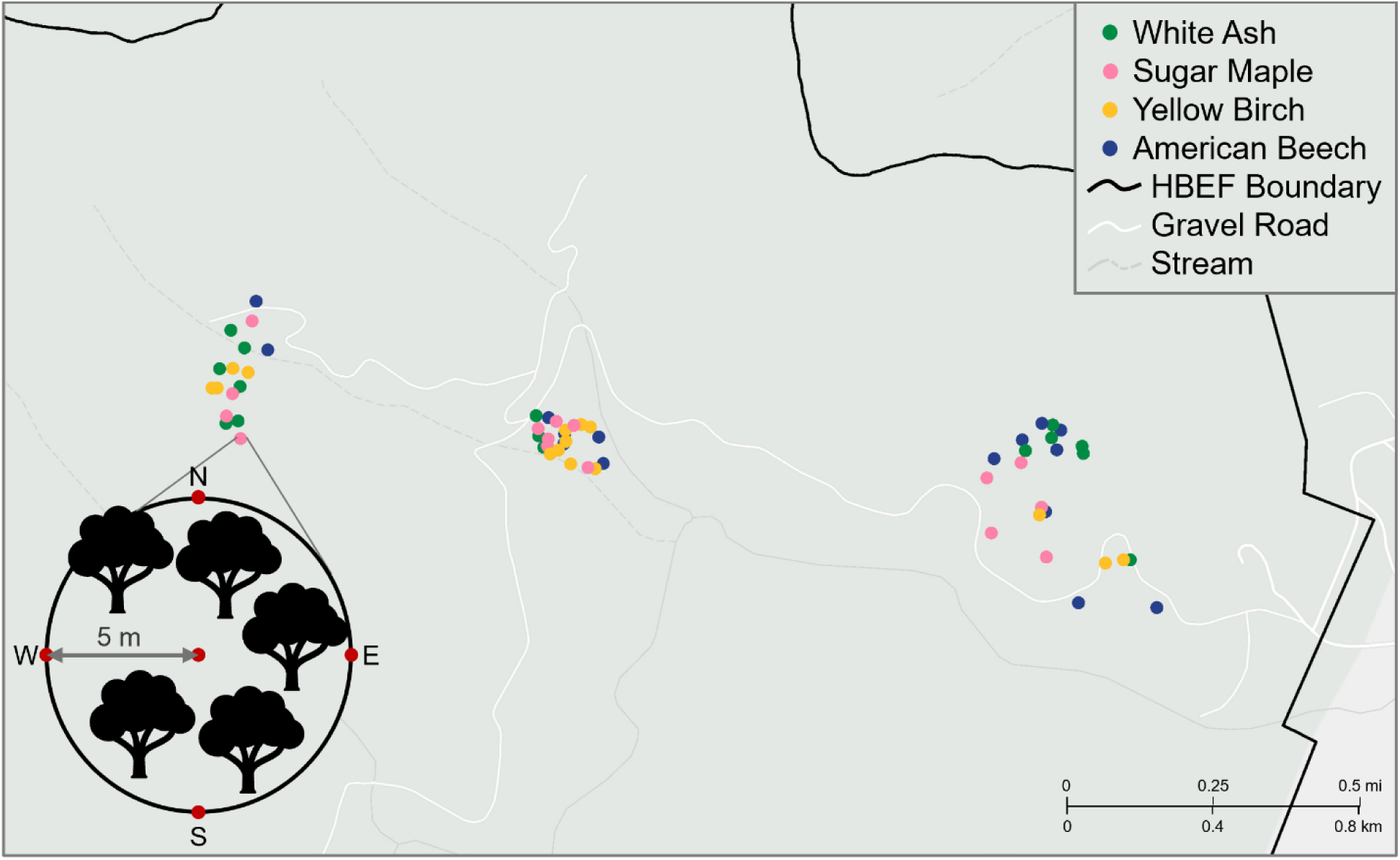
Map of the 60 study plots within the Hubbard Brook Experimental Forest, White Mountain National Forest, New Hampshire (43.95N, 71.75W). An example diagram of a plot’s layout is shown in the bottom left corner. All plots were at 203 to 609 m above sea level.

About half of the plots containing each focal trees species were situated on each of the two most common major soil types in HBEF: typical podzol or Bh podzol (Bailey et al., 2014). Typical podzols are characterized by the presence of a thin E horizon (mean 4.4 cm, Bailey et al., 2014) above one or two thicker spodic (Bhs, Bs) horizons. Bh podzols, formed by higher, fluctuating water tables, lack an E horizon and have a thick Bh horizon (mean 38 cm, Bailey et al., 2014). Plots were identified with the aid of hydropedological maps (Fraser et al., 2024) and soil types were verified via at least three soil pits (Studer et al., 2025).

Plots of all eight plot types were interspersed across three clusters which were separated by 1 – 2 km (Figure 1). Plots were 203 – 609 meters above sea level. Understory vegetation in the plots was dominated by a diverse assemblage of perennial herbs, along with ferns, bryophytes, tree saplings, and a few abundant shrubs (primarily *Viburnum lantanoides*) (Holmes and Likens, 2016b; Studer et al., 2026).

### Fungi Sampling

#### Belowground Sampling

Between July 28th and August 16th, 2022, five soil samples of the organic horizon were collected from each plot with a 2.54 cm diameter soil core. Each sample was frozen individually at -20° C within four hours of collection and stored for later DNA analysis.

In fall 2023, one or three soil samples from each plot were selected for DNA sequencing for a total of 143 samples. Five plots with each combination of tree and soil type were analyzed with three samples to enrich the data. Remaining plots were analyzed with one sample.

#### Aboveground Sporocarps

Two surveys of fungal fruiting bodies were conducted in 2023, the first in late July and the second two weeks later in early August. Few if any individual sporocarps persisted long enough to be counted in both surveys. For surveys, each circular plot was divided into eight equal pie-shaped subplots using the flagged cardinal directions. Sporocarps emerging from the soil were identified to genus and the number of individuals with a cap diameter of 2.5 cm or greater were counted for each sub-section. If the entirety of a cluster of cap-less taxa or connected small-capped mushrooms were longer than 2.5 cm they were also recorded. Sporocarps joined together above ground were counted as one individual but close-by individuals that were likely connected underground were still counted separately. If individuals were not identifiable to genus in the field, distinguishing characteristics were recorded, and a representative collected to make a spore print. Collected samples were dried and stored in paper bags for later reference.

### DNA Extraction and Bioinformatics

#### Illumina Sequencing

Fungal community DNA was extracted from soil samples using DNEasy Soil Pro Extraction Kits (Qiagen Science Inc.). Extraction followed the 2021 manufacturer’s protocol with a modification to the homogenization step. Samples of 250 mg of soil were homogenized for 40 seconds in a FastPrep-24 machine set at 6.0 m/s (MP Biomedicals). Large pieces of root and wood were removed ahead of extraction, but soils were not sieved to remove small roots or other fragments. DNA concentration in extracted samples was assessed using a NanoDrop One (Thermo Fisher Scientific) before proceeding to PCR amplification of the ITS1 region of ribosomal DNA. The forward primer ITS1-f (CTTGGTCATTTAGAGGAAGTAA) and reverse primer ITS2 (GCTGCGTTCTTCATCGATGC) were used in accordance with the Earth Microbiome Project (Thompson et al., 2017; Smith et al., 2018). PCR amplification was done with a Biometra TOne 96 G thermocycler (Analytik Jena) using KAPPA2G HotStart master mix (CustomBiotech) and the following cycling protocol: 95 °C/3 min; 30 repeats (95 °C/15 s, 52 °C/15 s, 72 °C/15 s); 72 °C/ 1 min; cooling to 12 °C. PCR products were cleaned using AMPure XP magnetic beads according to the manufacture’s instruction (Beckman Coulter) and DNA concentrations were quantified using a Qubit dsDNA BR Assay Kit (Thermo Fisher Scientific). Paired-end, 600 cycle sequencing was performed by Illumina NovaSeq2000 through the Genomics and Molecular Biology Shared Resource (GMBSR) at the Dartmouth Cancer Center (Hanover, NH). Returned sequences were processed using R software version 4.4.0 (R Core Team, 2024) following the DADA2 ITS-specific pipeline (Callahan et al., 2016). Unique operational taxonomic units (OTUs) were identified with the UNITE “eukaryotes 2” ITS database (Abarenkov et al., 2024a, 2024b) to the lowest confident taxonomic level using a naïve Bayesian classifier (DADA2 package).

#### Sanger Sequencing

Subsamples from 29 fungal sporocarps collected during field surveys were processed for DNA extraction following the same protocol used for the soil samples with two additions. Prior to homogenization, approximately 200 mg of each sporocarp samples were incubated at room temperature for 15 minutes in the DNEasy Soil Pro Extraction Kit (Qiagen Science Inc.) extraction buffer. Samples were then homogenized for 40 seconds in a FastPrep-24 machine set at 6.0 m/s, rested for 5 minutes, and homogenized a second time for 20 seconds at 6.0 m/s. DNA concentration in extracted samples was assessed using a NanoDrop One (Thermo Fisher Scientific) before proceeding to PCR amplification of both the ITS and RPB2 region of ribosomal DNA. A longer section of the ITS region and the addition of the RPB2 region were chosen for these samples as is recommended for achieving species level identification within highly conserved genera (Caboň et al., 2017; Li et al., 2019).

The forward primer ITS1-f (CTTGGTCATTTAGAGGAAGTAA) and reverse primer ITS4 (TCCTCCGCTTATTGATATGC) targeted the ITS region (White et al., 1990; Gardes and Bruns, 1993) while the forward primer bRPB2-6F (TGGGGYATGGTNTGYCCYGC) and reverse primer fRPB2-7cR (CCCATRGCYTGYTTMCCCATDGC) targeted the RPB2 region (Liu et al., 1999; Matheny, 2005). These ITS and RPB2 primers were added to separate subsamples of extracted DNA for each sporocarp resulting in 58 samples amplified by PCR. The PCR amplification procedure was the same as described for the soil samples except a different cycling protocol was used for the RPB2 region: 95 °C/3 min; 40 repeats (95 °C/15 s, 58 °C/15 s, 72 °C/15 s); 72 °C/ 1 min; cooling to 12 °C. PCR products were cleaned using AMPure XP magnetic beads and DNA concentrations were quantified using a Qubit dsDNA BR Assay Kit. Sanger sequencing was performed in the forward direction with an ABI 3730 Genetic Analyzer (Thermo Fisher Scientific) using ABI BigDye chemistry through GMBSR at the Dartmouth Cancer Center (Hanover, NH).

Returned sequences were trimmed by hand using sequence visualizations in FinchTV software version 1.4.0 (Geospiza Research Team, 2004) and identified to the lowest confident taxonomic level in “database nt” using the MegaBLAST program from the National Center for Biotechnology Information (Morgulis et al., 2008). ITS and RPB2 sequences were considered together for each sample to determine if species level identification was appropriate. Genus identity was accepted if our sample matched multiple database entries for the genus with a percent identity score > 97%. Identity information was then used to place previously unidentified sporocarps into the appropriate genus category within the sporocarp survey dataset.

### Data Filtration

Prior to statistical analysis, metabarcoding data were filtered as follows. First, we screened for likely contaminants using the “combined” method in the DECONTAM package (R software version 4.4.2), which assesses both the frequency of each OTU and the prevalence of each OTU in true positive versus negative control samples (Davis et al., 2018). One of the three negative controls did come back with OTU reads, which helped us identify 17 contaminant OTUs. The PHYLOSEQ package (McMurdie and Holmes, 2013) was then used to remove these contaminants, along with the negative controls, and failed positive samples with less than 100 total OTU reads. Following Kauserud (2023), we sought to reduce the over-splitting of genetically diverse fungal species into multiple OTUs by using the LULU R package (Frøslev et al., 2017) accompanied by VSEARCH (Rognes et al., 2016) to combine genetically similar OTUs into one OTU if they possessed the same distribution across the dataset. These corrections eliminated 1783 OTUs by combining them with appropriate “parent” OTUs. Next, we used PHYLOSEQ to remove OTUs that were not identified to a fungal phylum or had less than 5 reads total across all samples. We also removed one sample that had an unusually low read count of < 10000. These filtrations steps resulted in 135 curated samples representing 60 plots and containing 5588 ASVs. To reduce the stress in statistical analysis, we combined multiple samples from the same plot with simple addition to have one representative community per plot (10 plots were pooled for each of the four tree species). To account for bias from uneven library sizes, read counts were converted to relative percent abundance by dividing each of the 60 plots by their total read count using the “decostand” function in the VEGAN R package (Oksanen et al., 2024). We acknowledge that this normalization procedure can still result in elevated OTU richness in samples with higher read counts. Code for our filtering protocol, as well as our data, can be found on the Environmental Data Initiative (EDI) Portal (Ransom and Ayres, 2026).

### Soil Moisture Sampling

In each plot, we recorded soil moisture in the plot center and 0.5 meters from the plot edge in each cardinal direction. Moisture was measured as volumetric water content with the sensor needles inserted 5.5 cm into the ground (HOBO TEROS 12 US Soil Moisture/Temperature/EC sensor METER Group Inc.). Plots were sampled across two days and times (AM or PM) in August 2023, so sampling occasion was considered as an additional factor during analysis. All measurements were taken roughly 3 days after a significant rain event.

### Statistical Analysis

#### Fungal Community DNA

Soil fungal communities were characterized with respect to Shannon and Simpson diversity indices (VEGAN package). We compared diversity indices, observed OTU richness, and the percent of OTUs unique to each plot with a two-way ANOVA that included tree species, soil type, and their interaction as fixed effects (JMP Pro v17, SAS Institute, 2023). Post-hoc Tukey-Kramer HSD means comparisons were completed when appropriate (JMP Student Ed. v18, JMP Statistical Discovery, 2026). We also used the BIODIVERSITYR package (Kindt and Coe, 2005) to create rank abundance curves and compared OTU rank order.

We assessed our sampling effort and further compared the fungal communities by examination of taxonomic rarefaction functions. We used the R package iNEXT (Hsieh et al., 2016) with an input matrix of presence absence data from the 135 original samples before they were combined based on their plot label.

Patterns in fungal community structure were visualized and evaluated using a nonmetric multidimensional scaling (NMDS) ordination (Bray–Curtis distance and k = 2 axes; VEGAN package). We followed this with factorial analysis of tree species, soil type, and their interaction (PERMANOVA; “adonis2’’ function in VEGAN). These analyses were also performed with unrelativized data and with presence-absence data using a Jaccard distance matrix which each yielded similar results. Multivariate homogeneity of groups dispersions was then tested with the “betadisper” function in VEGAN. Given significant effects of tree species alone, we evaluated all pairwise comparison among species using the “pairwise.adonis” function which complements “adonis2” (Martinez Arbizu, 2020).

To explore which fungal OTUs were most strongly associated with different tree species, we searched for indicator species using the “multipatt” function in the INDICSPECIES package (Cáceres and Legendre, 2009). We also used the “strassoc” function within INDICSPECIES to identify which OTUs were only found under one tree species (A =1).

#### Sporocarp Surveys

Analysis of sporocarp survey data paralleled analysis of fungal community DNA. We analyzed the two sporocarp surveys separately for comparison. For each dataset, the “diversity” function in the VEGAN R package was used to calculate Shannon and Simpson diversity indices for each plot using count data. Diversity indices, total sporocarp count, mean sporocarp count, mean proportional abundance within a plot, genera richness, and the number of rare genera (< 10 sporocarps recorded in total) were evaluated with a two-way ANOVA that included tree species, soil type, and their interaction.

We calculated community dissimilarity matrices based on Bray-Curtis distance and fit NMDS ordinations with the VEGAN package (k = 2). Analyses used the proportion of eight sections of each plot occupied by a genus; these distributions were closer to normal and reduced ordination stress. Following Dexter et al. (2018), we compared our stress to a permutational null model of randomized genera abundances. The null model was produced with the quantitative swap and shuffle method “swsh_samp_r” in the “oecosimu” function in VEGAN which preserves fill and column and row frequencies, as well as row totals, and is appropriate for non-integer data. These models indicated that the ordination stress values were too high for reliable interpretations.

#### Soil Moisture

We analyzed soil moisture with an ANOVA model that included tree species, soil type, sampling occasion, and their interactions as fixed effects. Plot identity was treated as a random effect nested within tree species and soil type.

## Results

### Community DNA Taxa Distributions and Richness

We identified 5,588 unique fungal OTUs across our 60 study plots. More than half of these OTUs (58.8%) were only recorded in one study plot (Table 2). Three OTUs were found in all 60 plots, identified as *Saitozyma podzolica*, *Podila humilis*, and *Solicoccozyma terricola*. While almost half (43.9%) of OTUs could be matched to one of 505 distinct genera, 28.3% of OTUs were only identified to the level of fungal phylum. With > 100,000 reads total, the most common Ascomycete genera were *Geoglossum, Trichoderma, Oidiodendron*, and *Penicillium*; while the most common Basidiomycete genera were *Hygrocybe, Russula, Clavulinopsis, Saitozyma, Inocybe, Cortinarius, Amanita, Sebacina, Apiotrichum, Gliophorus, Solicoccozyma,* and *Ganoderma*. Two genera of Mortierellomycota, *Mortierella* and *Podila* also had > 100,000 reads total.

The greatest number of distinct fungal OTUs were found under white ash trees (Table 2, Supplemental Figure 1), although average OTU richness did not vary at the plot level (F(3,52) = 0.90, p = 0.45). There were significant differences in the prevalence of OTUs unique to a single plot (F(3,52) = 3.054, p = 0.036). At least 5% of OTUs detected in each plot were only found in that plot, with white ash plots supporting a higher percentage of unique OTUs (Figure 2). Sampling effort rarefaction models indicated increased sampling would continue to uncover many additional OTUs, with the most under ash (Supplemental Figure 1).

**Figure 2.**
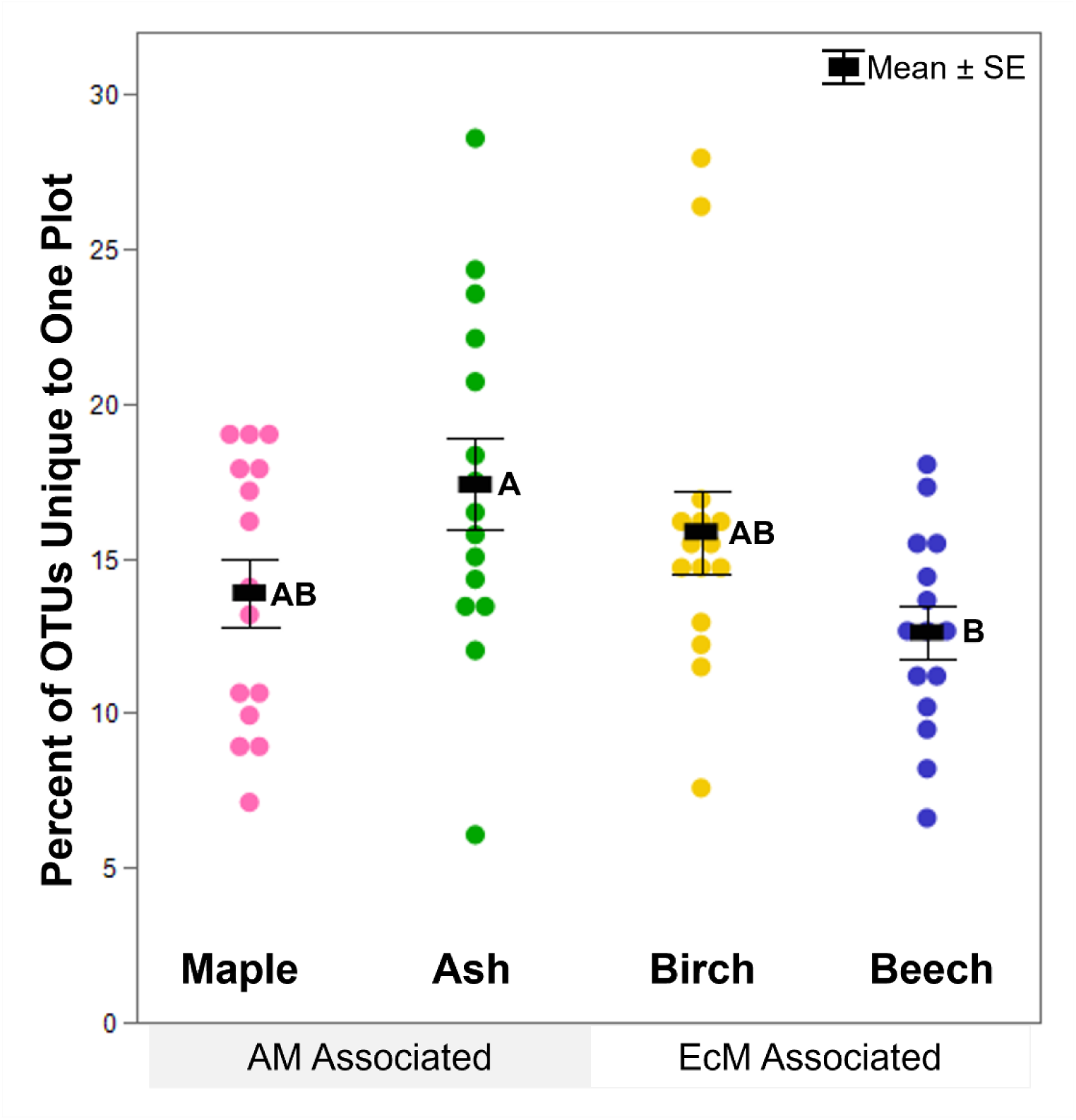
Percent of fungal OTUs found only in one plot varies according to tree species (F(3,52) = 3.05, p = 0.036). Points represent each of the 60 study plots (N = 15 for each tree species). Letters correspond to post-hoc Tukey HSD means comparison.

There were no differences in Shannon or Simpson diversity indices (F(3,52) = 0.39, p = 0.76 ; F(3,52) = 1.19, p = 0.32), which affirmed Rank Abundance Curves (Figure 3). Rank abundance curves show that the average fungal OTU community under each tree was composed of over 100 taxa, with the majority of those taxa individually contributing less than 1% to the community’s total abundance. While patterns in community richness and evenness were similar across all trees, visualizing species rank according to OTU identity revealed differences in the most abundant OTUs under each tree (Figure 4). None of the most abundant OTUs were among the top 10 most common under all four trees.

**Figure 3.**
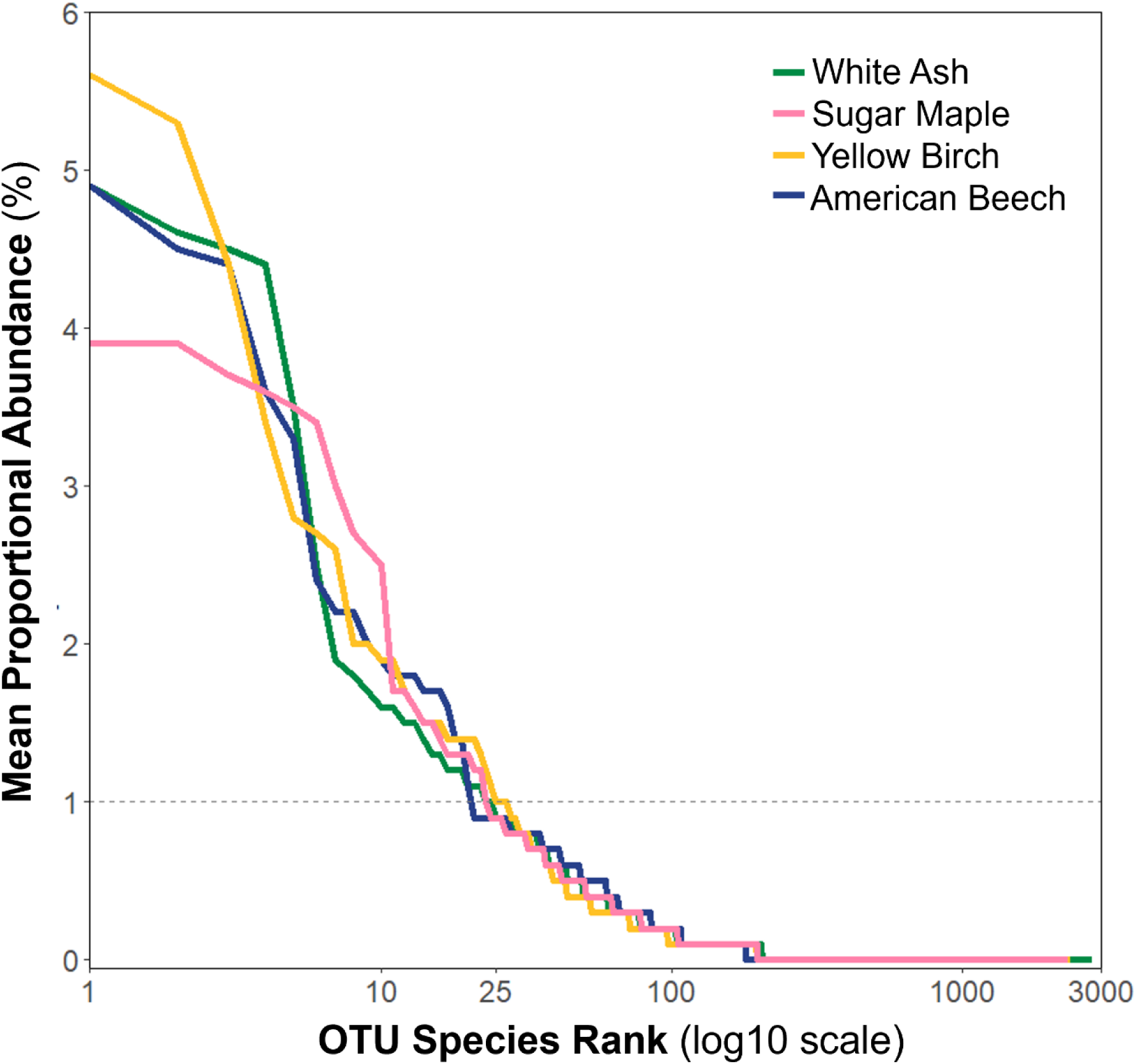
Fungal OTU Rank Abundance Curves depict similar patterns in fungal community richness (curve length) and evenness (curve slope) across tree species.

**Figure 4.**
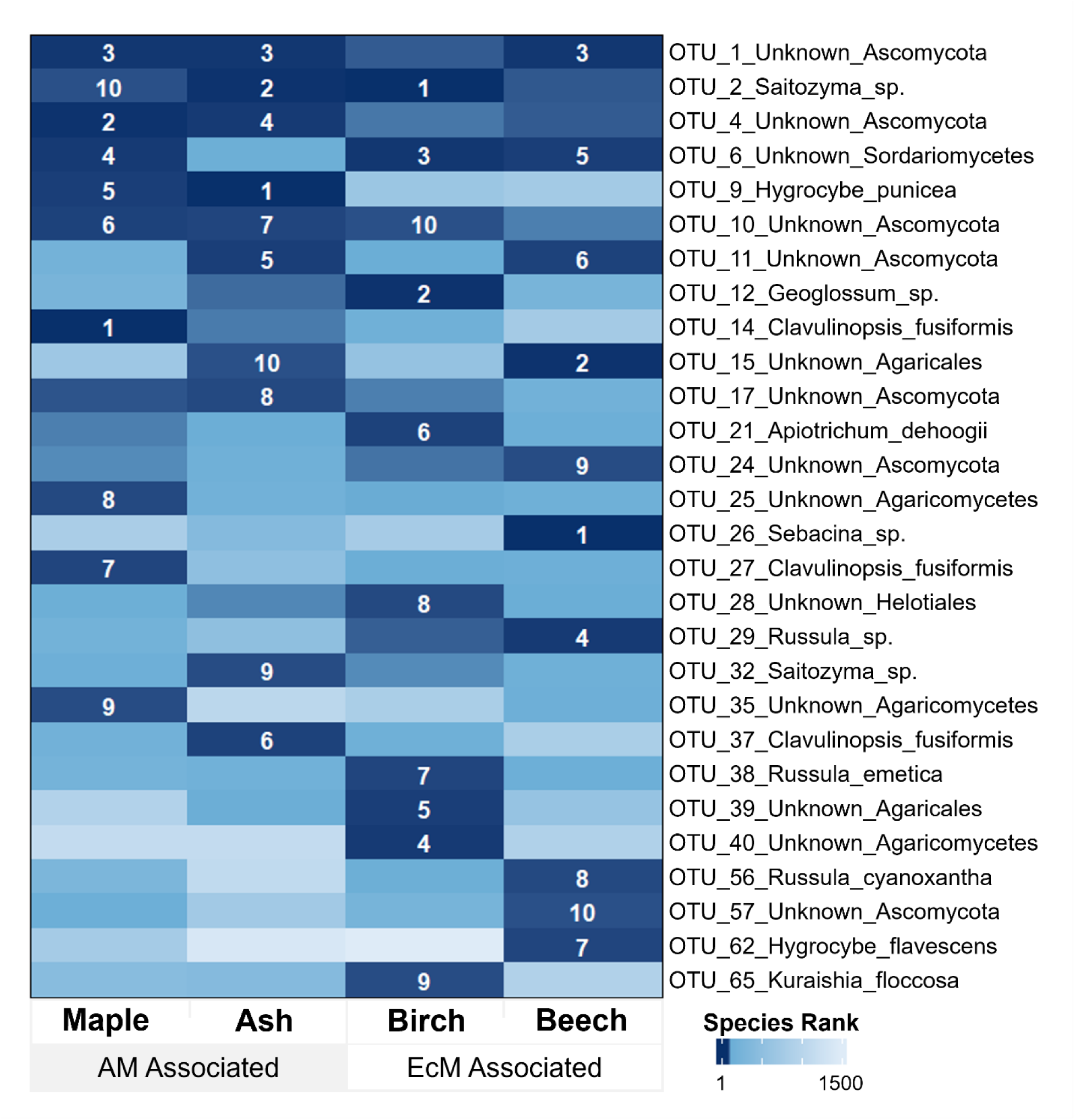
Identity of the most abundant fungal OTUs differed across tree species. Displayed OTUs are the top 10 most abundant taxa under at least one tree species. Heatmap colors represent species rank values, with darker colors indicating higher mean proportional abundance. Species rank values up to 10 are indicated in white.

Soil type and the interaction between tree and soil type had no significant association with richness or diversity indices at any level.

### DNA Community Patterns

Soil samples showed distinct fungal community structure based on tree species (Figure 5, k = 2, stress = 0.189). Two dimensions (k=2) were deemed appropriate through visual interpretation of a scree plot of ordination stress (stress value > 0.2; Supplemental Figure 2). Tree species identity, but not soil type showed pattern in the ordinated community structure (F(3,52) = 1.79, p = 0.001, Table 1A). Checks on beta dispersion confirmed that within-group variation among tree species did not significantly influence these results (F(3,56) = 0.66, p = 0.58).

**Figure 5.**
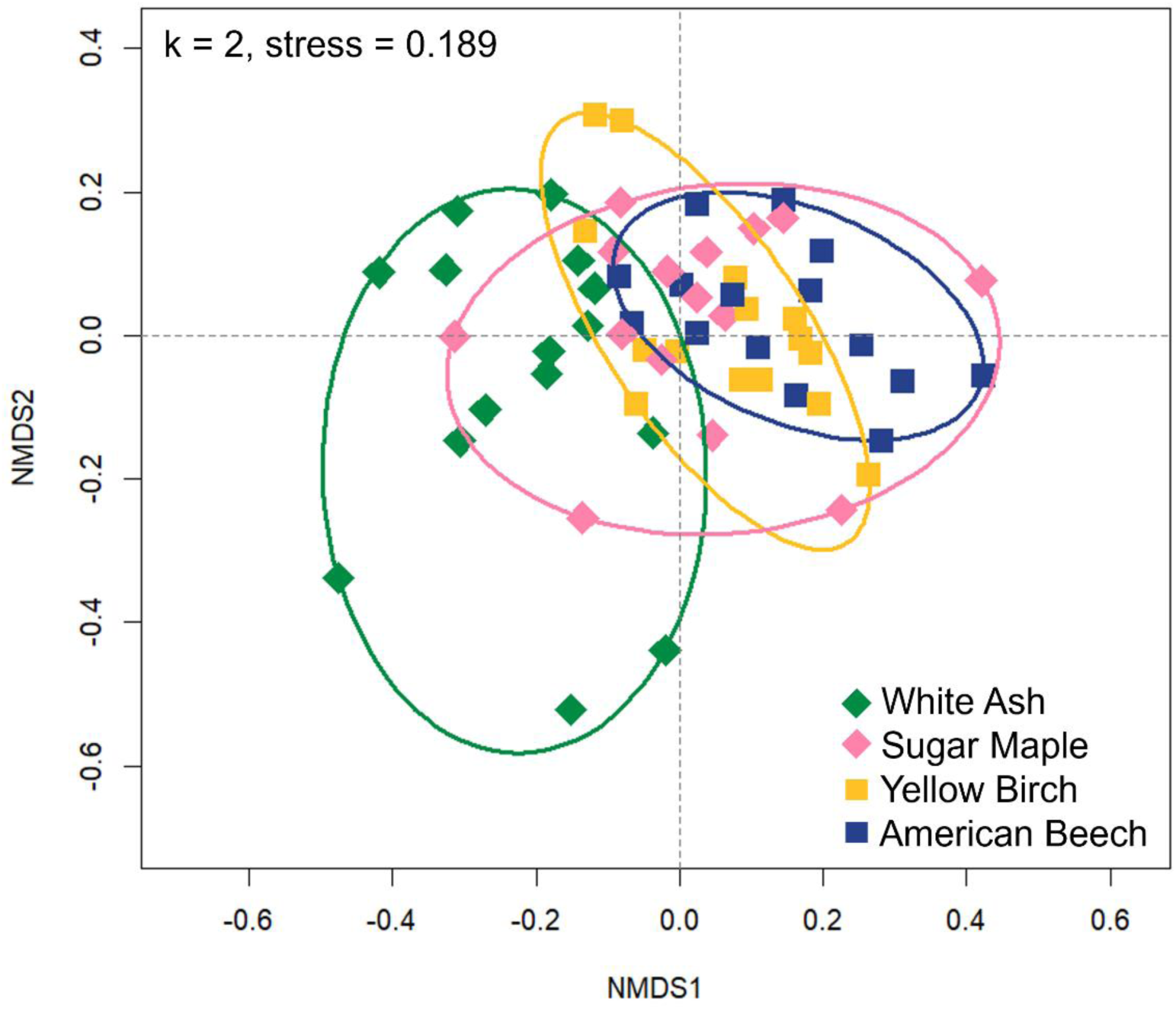
Only plot tree species reflects a significant pattern in fungal OTU community structure (F(3,52) = 1.79, p = 0.001). Nonmetric multidimensional scaling ordination with ellipses enclosing the 15 replicate plots of each tree species (total N= 60).

**Table 1.**
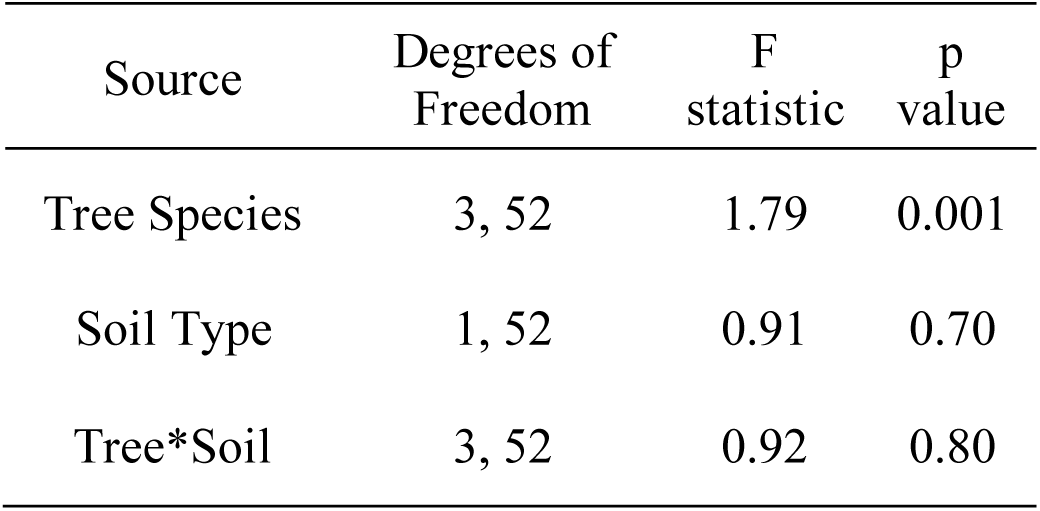
A. Two-way PERMANOVA results comparing fungal community composition across tree species, soil type, and their cross interaction.

**Table 1.**
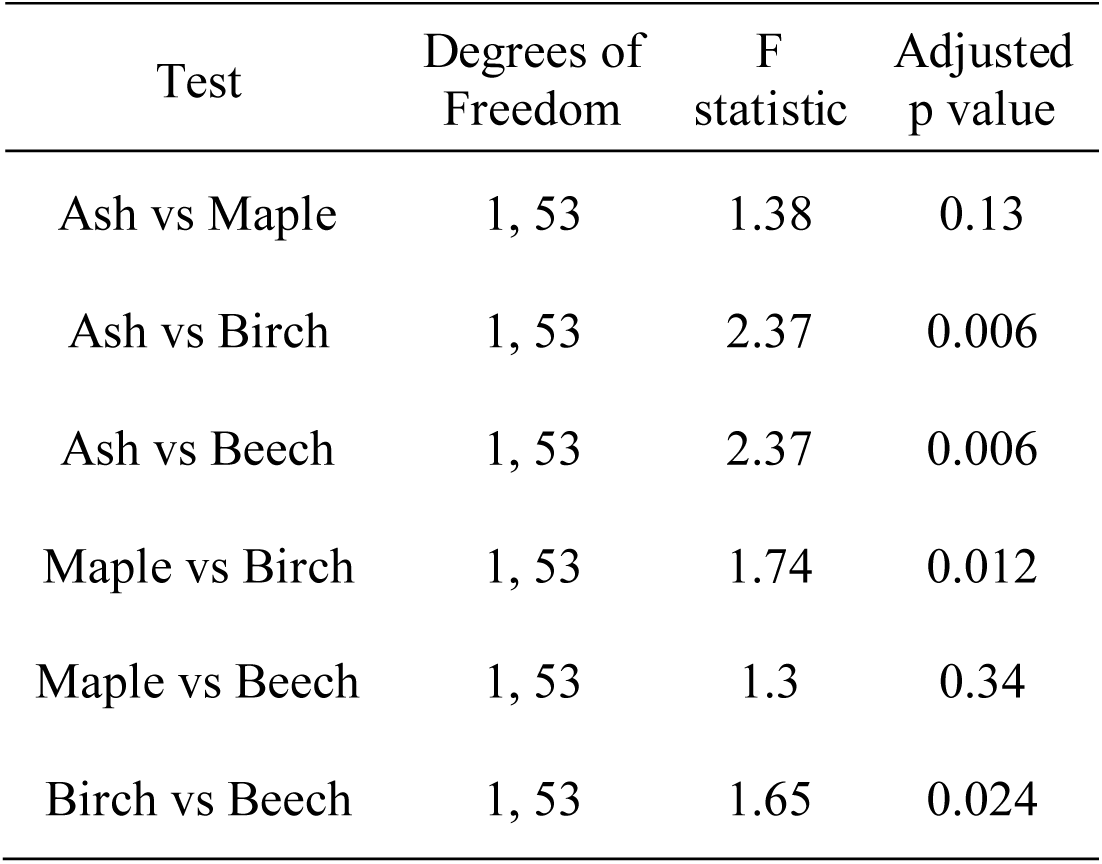
B. Pairwise PERMANOVA results testing differences between soil fungal communities beneath the four dominant tree species: white ash, sugar maple, yellow birch, and American beech.

Visual interpretation of the ordinated fungal community suggested white ash trees differed from the other species present but were most similar to sugar maple. Subsequent pairwise PERMANOVA comparisons confirmed that the fungal communities under ash trees differed significantly from the EcM associated beech and birch trees (Table 1B). Two out of the fifteen sugar maple plots had fungal communities that were especially similar to ash (Figure 5), driving the lack of statistical differences between ash and maple communities overall (F(1,53) = 1.38, p = 0.13, Table 1B). Surprisingly, the fungal community under sugar maples, an AM associated tree, were not significantly different from those under the EcM associated American beech (F(1,53) = 1.30, p = 0.34, Table 1B). Additionally, the two EcM associated trees, American beech and yellow birch, had different fungal communities despite the partial overlap of their community ellipses (F(1,53) = 1.65, p = 0.024, Figure 5, Table 1B).

### Indicator Species Analysis (ISA) Across Tree Types

Strikingly, we identified 90 indicator OTUs for white ash plots, with roughly 30 indicator OTUs for each of the other tree species (Table 2, Supplemental Tables 2A-D). All indicator OTUs had significant p values ranging from p = 0.001 to p = 0.05 (Supplemental Tables 2A-D). Indicator OTUs included 52 fungal genera, but 61% lacked genus level taxonomic information, and 32% were only identified to phylum. The majority of indicator OTUs were Ascomycetes (75%) with most others matching Basidiomycetes.

**Table 2.** Summary table characterizing the community of fungal OTUs summed for each tree species.

| Tree Species | Plots | Total OTUs | OTUs Unique to One Plot | OTUs Unique to Tree Species <sup>a</sup> | Indicator OTUs <sup>b</sup> |
| --- | --- | --- | --- | --- | --- |
| Sugar Maple | 15 | 2315 | 752 | 48 | 26 |
| White Ash | 15 | 2790 | 1085 | 144 | 90 |
| Yellow Birch | 15 | 2364 | 859 | 44 | 37 |
| American Beech | 15 | 1960 | 591 | 28 | 33 |
| Total | 60 | 5588 | 3287 | 264 | 186 |

a. OTUs Unique to Tree Species were found in multiple plots of one tree species. Some of these OTUs were also identified as Indicator OTUs.
b. Indicator OTUs determined through Indicator Species Analysis, see supplemental tables 2A-D for details including p values.

We also identified 264 OTUs found in multiple plots but only under one tree species (Table 2.) Of these OTUs unique to tree species, 54.54% were found only under white ash. Only some of these OTUs unique to a tree species were statistically considered indicator OTUs.

Most indicator OTUs (87%) were found at least once under a tree species other than the one they were significant indicators for. On average, an indicator OTU was found in 45% of the plots of its tree species, and only 5 indicator OTUs were found in every plot of their tree species (B mean = 0.447, SD = 0.202, min = 0.20, max = 1).

### Sporocarp Survey Distributions and Analysis

Across the two surveys, we recorded 1667 sporocarps in 37 genera plus two additional family level groups (Table 3). Members of the Hygrophoraceae and Boletaceae families could not be easily identified to genus, with the exception of boletes within *Strobilomyces* which were counted separately. The most commonly observed sporocarps were within *Russula* (338 observations) with members of the Hygrophoraceae family and the *Gymnopus* and *Coltricia* genera also observed over 100 times. The genera *Amanita, Craterellus, Cortinarius, Entoloma, Lactarius,* and *Leotia*, along with the Boletaceae family each had more than 50 observations (Figure 6). Other taxa were less common, with 16 ‘rare’ genera observed > 10 times (Table 3).

**Figure 6.**
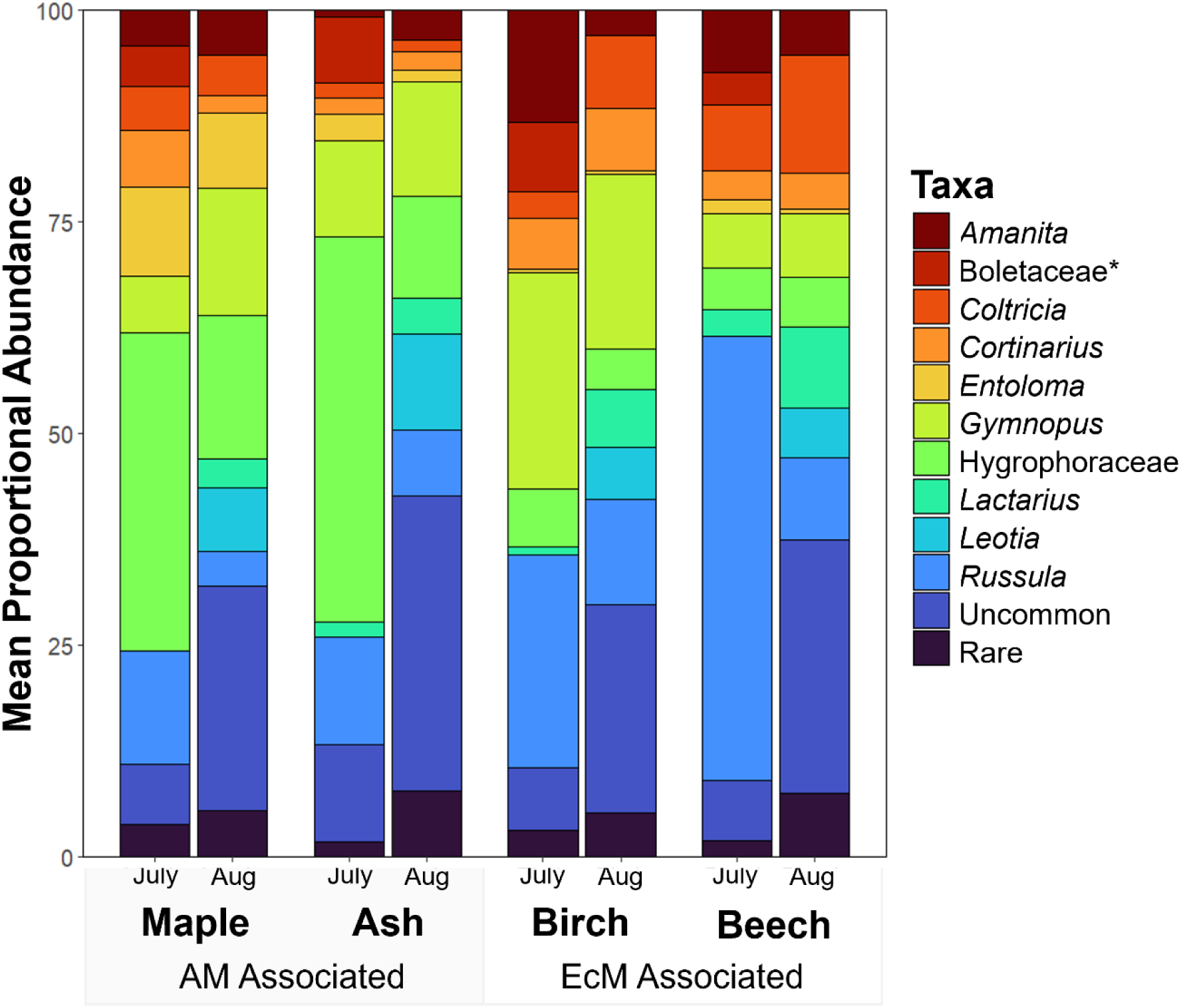
Proportional abundance of sporocarp taxa under four tree species, as observed during two survey periods. ‘Rare’ taxa were observed < 10 times across both surveys. ‘Uncommon’ taxa were observed < 50 but > 10 times. *Taxa in the *Strobilomyces* genus were counted separately from other members of the Boletaceae family.

**Table 3.**
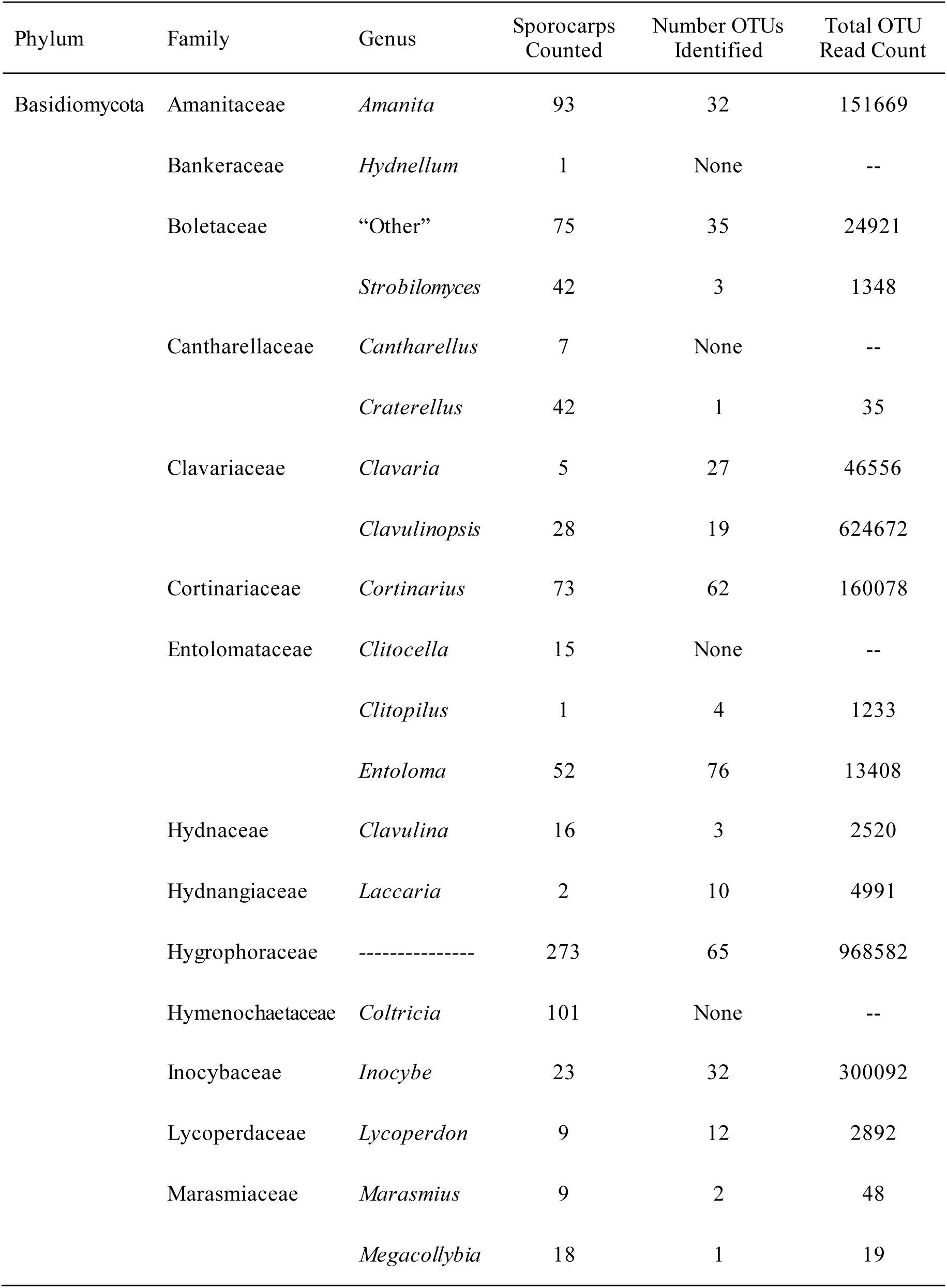
A. Fungal taxonomic groups identified during both rounds of sporocarp surveys compared to corresponding OTUs identified from soil eDNA.

**Table 3.**
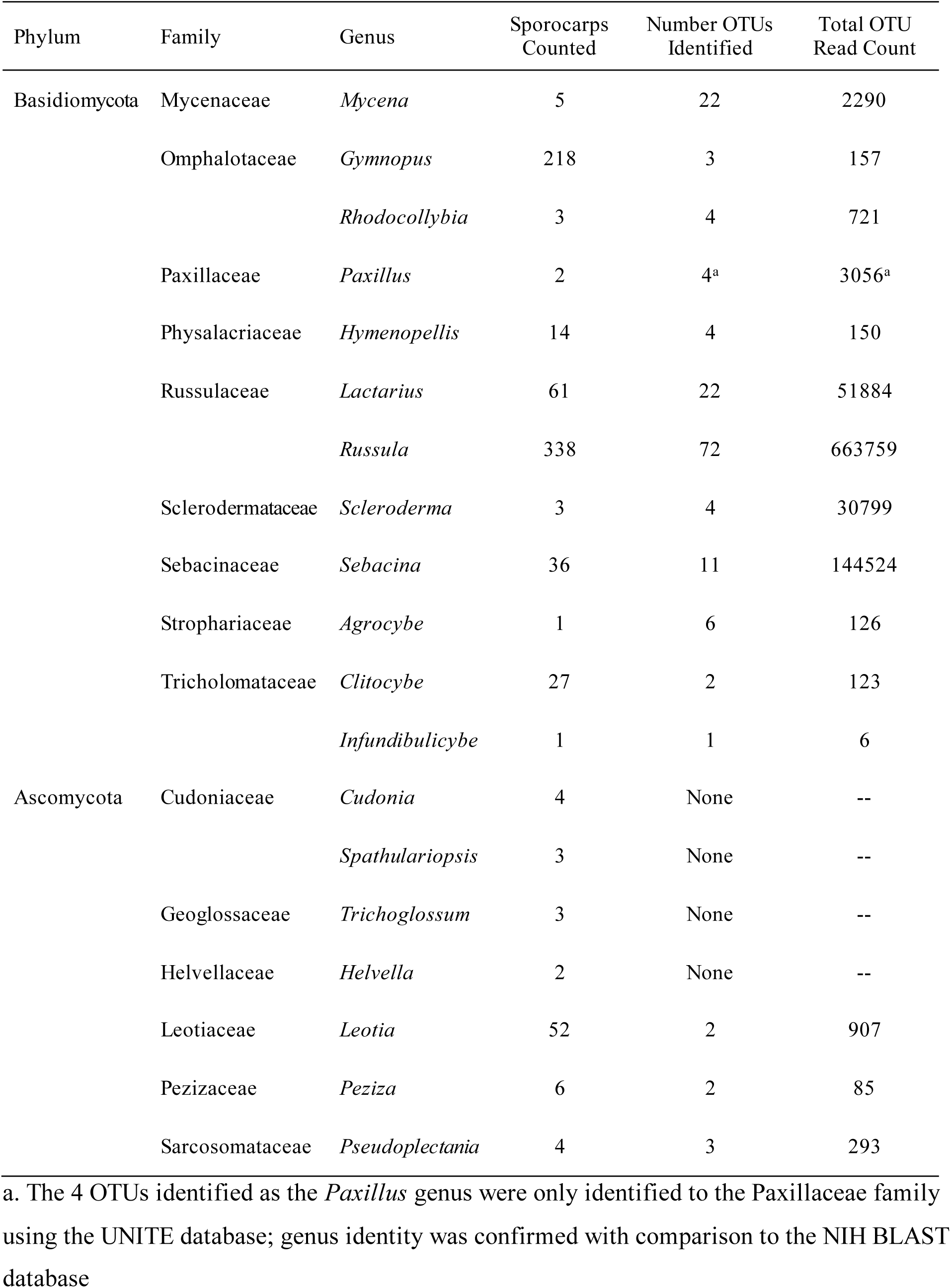
B. table continued from previous page.

ANOVAs revealed no significant differences across study plots in sporocarp abundance, taxonomic richness, or either of the two diversity indices. Proportional data from surveys 1 and 2 were also ordinated as Bray-Curtis distance matrices (k = 2, S1 stress = 0.22, S2 stress = 0.23), but biological interpretation was deemed inappropriate based on permutational null models (S1 mean simulated stress = 0.23, Z = - 0.17, p = 0.28; S2 mean simulated stress = 0.22 Z = -0.42, p = 0.58.). Still, graphs of proportional abundance separated by tree species revealed interesting patterns in sporocarp communities among trees and over time (Figure 6).

### Comparison of Community DNA and Sporocarp Data

There were numerous similarities between data from sporocarp surveys and community DNA. *Russula* was the most frequently observed taxon in sporocarp surveys and the second most common Basidiomycete found in soil samples (total OTU read count, Table 3). The first most common Basidiomycete genus in soil samples, *Hygrocybe*, along with the common *Gliphorus* genus, belong to the second most observed taxa group in our aboveground surveys: the Hygrophoraceae family. The genera *Amanita, Cortinarius*, and *Entoloma* were also among the most recorded genera in both data sets.

There were also differences between our two fungal community datasets. Even though the *Gymnopus* genus was one of the top four taxa groups observed in sporocarp surveys, *Gymnopus* had a total of only 157 reads across 3 OTUs in the community DNA data (Table 3). The genus *Coltricia*, fourth most common taxon in sporocarp surveys, was never detected from DNA. Seven other genera were identified through visual characteristics in our sporocarp surveys but were not matched to any OTUs in our community DNA data: *Hydnellum, Cantharellus, Clitocella, Cudonia, Spathulariopsis, Trichoglossum*, and *Helvella* (Table 3). Of these missing genera, the most observed was *Clitocella* which was recorded 15 times.

### Soil Moisture

Soil moisture differed across tree species and soil type with no significant interactions (Supplemental Figure 3, Supplemental Table 1). As expected, Bh podzol soils were more moist (least square mean ± 1SE = 0.335 ± 0.009, F (1,36) = 14.74, p = 0.0005) than typical podzols (LSqM ± 1SE = 0.284 ± 0.009), although a wide range of moisture levels were recorded in each soil type. Soils under white ash trees tended to be more moist (LSqM ± 1SE = 0.354 ± 0.013) than soils under other tree species: LSqM ± 1SE for beech, maple, and birch = 0.311 ± 0.014, 0.285 ± 0.013, and 0.287 ± 0.013, respectively.

## Discussion

DNA metabarcoding of soil samples revealed differences in fungal community structure beneath the four tree species included in this study. These differences were unrelated to soil type and were not simply explained by tree mycorrhizal association. White ash trees stood out with a particularly distinct fungal community. Across ash plots, there were more fungal OTUs unique to a single plot, more OTUs found only under ash, and more than double the number of significant indicator OTUs compared to other trees. NMDS ordination showed white ash fungal communities were different from those of American beech and yellow birch and were also largely separated from sugar maple plots (Figure 5). Fungal communities under American beech and yellow birch trees were significantly different from each other, despite their shared symbiosis with EcM fungi. Ultimately, our results suggest that species-specific features of canopy trees may be important to soil fungal communities. We add to the growing body of work uncovering differences in fungal communities below trees with the same mycorrhizal type (van der Linde et al., 2018; Lang et al., 2021; Hicks Pries et al., 2023).

Our findings of distinct fungal communities under white ash confirm the risk of fungal co-extinction with the impending extirpation of ash trees. The 90 indicator OTUs associated with ash, the 144 OTUs that were only found under ash, and especially the 16 OTUs in both categories can be considered at risk for co-extinction (Table 4). Differences observed in fungal communities across larger scales (Talbot et al., 2014; van der Linde et al., 2018; Moore et al., 2021) necessitate study of ash-associated fungal communities in other forests to confirm vulnerability of specific fungi. While many mycologists assert that fungi functional groups are more important to ecosystems than species identity (Bödeker et al., 2016; Zanne et al., 2020; Anthony et al., 2022), it is challenging to assess the functional roles of indicator OTUs without genus level identification (Nguyen et al., 2015). Even for known species, functional roles can shift (Zanne et al., 2020) or remain poorly characterized (Põlme et al., 2020; Tedersoo et al., 2022). Regardless of which species are at risk, a decline in the genetic diversity of forest fungi may compromise the abilities of these communities to adjust to future environmental challenges (Nic Lughadha et al., 2020).

**Table 4.** Fungal OTUs at risk of coextinction due to extinction of ash. These taxa were found only in ash plots and were also identified as ash indicators by Indicator Species Analysis (ISA)

| Phylum | OTU Label | Taxon Name | Total OTU Read Count | Number of Plots | Indicator Value | p Value |
| --- | --- | --- | --- | --- | --- | --- |
| Ascomycota | OTU_5481 | *Alternaria* sp. | 25 | 5 | 0.577 | 0.004 |
|  | OTU_3623 | *Brachysporium* sp. | 87 | 4 | 0.516 | 0.011 |
|  | OTU_2845 | *Gliocladium* sp. | 150 | 4 | 0.516 | 0.009 |
|  | OTU_3674 | *Ochrocladosporium frigidarii* | 84 | 5 | 0.577 | 0.005 |
|  | OTU_1746 | *Pleotrichocladium opacum* | 427 | 4 | 0.516 | 0.004 |
|  | OTU_3029 | *Spirosphaera* sp. | 132 | 4 | 0.516 | 0.012 |
|  | OTU_2525 | *Tolypocladium* sp. | 198 | 4 | 0.516 | 0.011 |
|  | OTU_2082 | *Trichoderma polysporum* | 313 | 4 | 0.516 | 0.012 |
|  | OTU_1153 | Unknown Ascomycota | 1096 | 5 | 0.577 | 0.002 |
|  | OTU_627 | Unknown Ascomycota | 2939 | 4 | 0.516 | 0.012 |
|  | OTU_1355 | Unknown Ascomycota | 708 | 4 | 0.516 | 0.015 |
|  | OTU_2166 | Unknown Ascomycota | 275 | 4 | 0.516 | 0.014 |
|  | OTU_2453 | Unknown Ascomycota | 211 | 4 | 0.516 | 0.008 |
|  | OTU_2990 | Unknown Ascomycota | 136 | 4 | 0.516 | 0.014 |
|  | OTU_2991 | Unknown Dothideomycetes | 136 | 4 | 0.516 | 0.015 |
| Basidiomycota | OTU_4606 | Unknown Tremellodendropsidales | 44 | 3 | 0.447 | 0.044 |

While ash specialists were of particular concern to us, common fungi also contributed to the observed differences in communities among trees. Rank abundance curves showed that the average fungal community was dominated by around 25 OTUs, with many additional rare taxa, regardless of tree type (Figure 3). However, the identity of these dominant OTUs differed among trees, with 21 out of 28 OTUs only falling within the top 10 most common taxa for one tree (Figure 4). These figures indicate that differences in the rank value of common OTUs are contributing to the divergence of the fungal communities under different trees. Reordering, or changes in the relative abundance of species, is one of the fundamental mechanisms of community change that can greatly alter community dynamics (Collins et al., 2008; Avolio et al., 2019, 2021). Observed differences in fungal OTU order among trees suggest that the loss of ash will lead to species reordering in forest fungal communities.

Analyses of our sporocarp survey data were limited by taxonomic resolution but support several interesting patterns. Even at the genus level, there was evidence of temporal shifts in sporocarp communities between the two surveys (Figure 6). This aligns with findings of other sporocarp studies with species-level identification, which confirm long-observed variations in fungal fruiting phenology (Eveling et al., 1990; Sato et al., 2012; Brown et al., 2022). These differences appear to be driven by plant phenology (Dickie et al., 2010; Sato et al., 2012) and other environmental variables (Sato et al., 2012; Bashian-Victoroff et al., 2025). While biological interpretation of NMDS ordinations for our sporocarp data was inappropriate, comparing sporocarp abundance among tree species suggests these communities could be distinct (Figure 6). In particular, future sporocarp studies with species-level resolution might want to investigate how *Entoloma*, *Gymnopus*, Hygrophoraceae, and *Russula*, species vary with tree species. Continued research on fungal phenology and sporocarp communities will be essential to understanding how fungi are being impacted by global change.

Comparison between our soil metabarcoding and sporocarp survey datasets highlighted informative similarities and differences. Most genera that were frequent as sporocarps were also detected frequently in community DNA samples (Table 3). Our sporocarp surveys confirmed that known EcM genera such as *Russula*, *Cortinarius*, and *Amanita*, are indeed living and reproducing under AM associated ash and maple trees. Additionally, if OTUs per genus approximates the true species richness, the DNA suggests, for example, that there are around 70 species of *Russula* within HBEF. This seems plausible based on field observations when conducting our sporocarp surveys. Discrepancies between our data sets illustrate other known limitations in sampling and interpreting fungal community DNA (Kauserud, 2023). Though the ITS1 region is preferred for sequencing many fungi (Thompson et al., 2017; Li et al., 2019), it poorly captures certain groups, including the *Cantherellus* genus (Schoch et al., 2012), and potentially the *Coltricia* genus (Tedersoo et al., 2007). Indeed, *Cantherellus* and *Coltricia* were found in our sporocarp surveys but not in community DNA. Alternatively, *Coltricia* (ectomycorrhizal but sometimes wood associating (Tedersoo et al., 2007)) and other taxa, including *Gymnopus* (leaf litter saprotroph (Põlme et al., 2020)), may be underrepresented in DNA samples due to a mismatch between their habitat preferences and the soils we sampled. Further investigations are needed to better understand relations among DNA metabarcoding, soil hyphae, mycorrhizal root communities, and sporocarps.

The discrepancies we observed among fungal communities under different tree species were likely influenced by species-specific tree traits. One attribute that might influence soil fungi is the lability of leaf litter, which varies among tree species and influences physical and chemical soil conditions (Jacob et al., 2009; Heděnec et al., 2023). AM associated tree leaves generally decompose faster than EcM associated leaves due to lower C:N ratios and reduced lignin content (Melillo et al., 1982; Fernandez and Kennedy, 2016). However, important variation in leaf chemistry exists even among species the same mycorrhizal type (Phillips et al., 2013). Measurements of leaf chemistry in the same study forest (Lang et al. 2021), showed that white ash and yellow birch leaves have similarly low C:N ratios relative to sugar maple and American beech. This is consistent with differences in fungal communities between birch and beech trees, and with similarities between beech and maple. Surprisingly, Studer et al. (2025) did not find lower C:N ratios in the combined organic soil horizons of the same birch plots used in this study, indicating that leaf traits do not necessarily translate to long term differences in soil chemistry. Other species-specific traits, such as the production of root exudates, may also contribute to unique soil communities (Hupperts et al., 2017; Vives-Peris et al., 2020; Keller et al., 2021), but further research is needed to better understand how such cryptic traits vary among trees.

Another possible driver of differences in fungal communities is variation in animal-mediated fungal dispersal. Even among wind-dispersed fungi, animal interactions can influence both spore transport and establishment success (Elliott et al., 2019; Stephens and Rowe, 2020; Elliott et al., 2022; Borgmann-Winter et al., 2023). Tree traits or other environmental differences associated with the tree species in our study could influence animal activity leading to localized differences in fungal dispersal within a forest. White ash groves in HBEF are also distinguishable in having more abundant and diverse understory vegetation (Studer et al., 2026) In addition to direct effects, understory vegetation may influence animal activity, which in turn shapes fungal communities (Cázares and Trappe, 1994; Dundas et al., 2018; Borgmann-Winter et al., 2023). Microhabitat selection by small mammals (Miller and Getz, 1977; Kaminski et al., 2007; Nelson et al., 2019), and birds (Holmes and Robinson, 1988; Korňan et al., 2013), is influenced by understory vegetation. *Trillium erectum*, a perennial flowering herb browsed by white-tailed deer (Vellend et al., 2003), is more common under ash trees, and white-tailed deer are competent dispersers of fungal propagules (Cadotte et al., 2021; Elliott et al., 2022). If animals are preferentially foraging or utilizing the understory in other ways, they are likely shaping the fungal communities under ash as they move spores throughout the forest.

While we did not see patterns in fungal community associated with hydropedological soil types, we do not discount the possibility of edaphic factors influencing soil fungi. The two soil types in our study are both podzols and similar with respect to some traits that might matter to fungi. Studer et al. (2025) found the two soil types to be similar in soil pH and %N. Soil pH, soil N, and soil moisture have frequently been linked to distinct fungal communities (Talbot et al., 2014; Taylor et al., 2014; van der Linde et al., 2018; Steidinger et al., 2020). Our soil moisture measurements identified Bh podzols as wetter than typical podzols, but with high within-plot variation, differences in mean soil volumetric water content did not translate to fungal communities. Still, soil moisture might play a role in differentiating fungal communities among tree species, as white ash plots had higher average soil water content. It is also possible that differences among soil types would emerge from sequencing individual mycorrhizal root tips, rather than DNA from the soil. *Cenococcum* species were detected occasionally from soils in our study, but Vietorisz (2019) visually identified *Cenococcum geophilum* on more than 85% of 6200 EcM-colonized root tips collected within HBEF. This discrepancy may be explained by *C. geophilum*’s resistance to decomposition or seasonal growth patterns (Fernandez et al., 2013), but further study of root tips may confirm distinct community patterns. Nevertheless, evidence from community DNA and sporocarps indicates that the two hydropedological soil types in this study have quite similar communities of soil fungi once the effects of canopy tree species have been accounted for.

Overall, our results indicate that white ash support a different community of soil fungi compared to the three other deciduous tree species studied. The regional community of soil fungi in forests of the White Mountains will likely change due to the oncoming functional extinction of ash and possible fungal coextinction. Ultimately, the future composition of forest fungi will depend in part on which tree species replace ash. This succession is challenging to predict, as changing climate and other anthropogenic impacts are simultaneously shaping the future of northeastern forests. On one hand, forests tend to shift towards AM tree dominance as they age and nitrogen deposition may favor AM trees that can more easily access the additional inorganic nutrients (Jo et al., 2019; Moore et al., 2021). On the other hand, Jo et al. (2019) found that forests in the New England “warm continental” ecoregion currently have more EcM tree saplings than AM tree saplings in the understory, suggesting future EcM tree dominance in this region. Increasingly warm and wet conditions may also permit the colonization of regional forests by tree species for which the White Mountains are beyond their current distributions (Prasad et al., 2020). Continuing long-term studies will be crucial for understanding the effects of tree species loss on the structure and function of forest ecosystems.

## Acknowledgments

This project was supported by National Science Foundation funding for Long Term Ecological Research at the Hubbard Brook Experimental Forest (Grants 1637685 and 2224545) and the Kaminsky Family Fund awarded through the Dartmouth Undergraduate Research Programs to FRR. Metabarcoding analysis was made possible by the National Cancer Institute Cancer Center Support Grant 5P30 CA023108-41 which supports the GMBSR Shared Resources facility at the Dartmouth Cancer Center. Many thanks to K.L. Cottingham for her guidance and editing and to the anonymous reviewers who strengthened this manuscript. The authors have no conflict of interest to declare.

## CRediT Author Contributions

**Farrar R. Ransom:** Conceptualization, Methodology, Formal analysis, Investigation, Data Curation, Writing - Original Draft, Writing - Review & Editing

**Paul Metzler:** Methodology, Formal analysis, Data Curation, Writing - Review & Editing

**Elizabeth A. Studer:** Methodology, Writing - Review & Editing

**Matthew P. Ayres:** Conceptualization, Methodology, Formal analysis, Writing - Review & Editing

**V. Bala Chaudhary:** Conceptualization, Methodology, Resources, Writing - Review & Editing

