## Supplemental Tables and Figures for "Unique soil fungal communities are associated with disappearing ash trees in a northern temperate hardwood forest"

**Supplemental Figures**

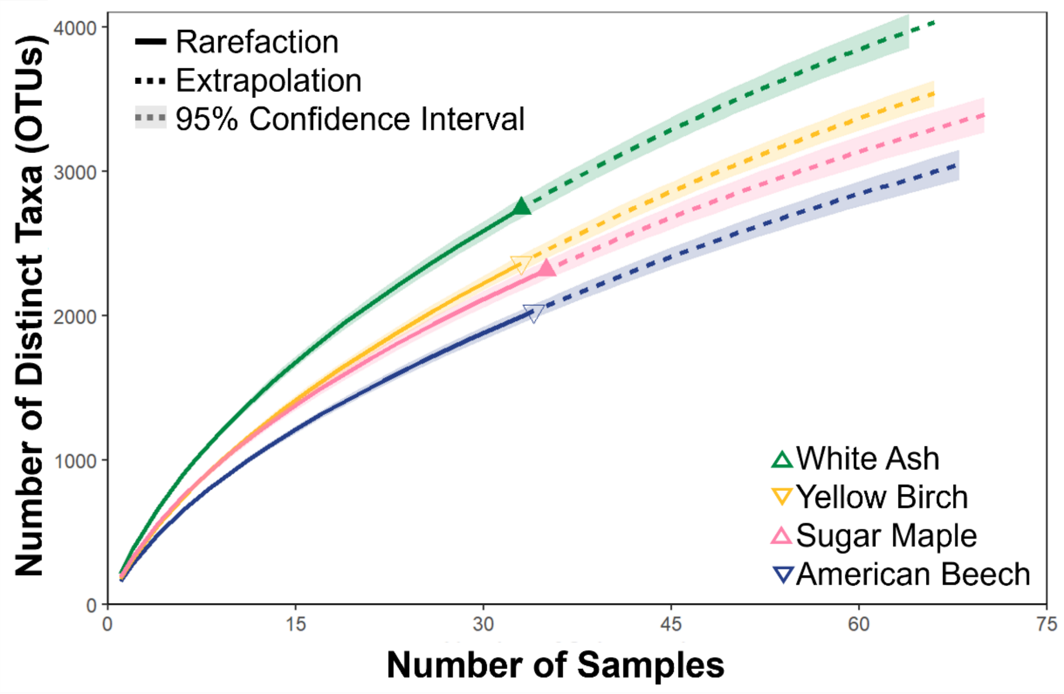


**Supplemental Figure 1.** Sampling effort curves reveal differences in species richness among tree species. The number of samples for each tree species were N = 33 for ash and birch, N = 35 for maple and N = 34 for beech. Samples taken from the same plot were combined for other analysis, see data filtration section of methods.

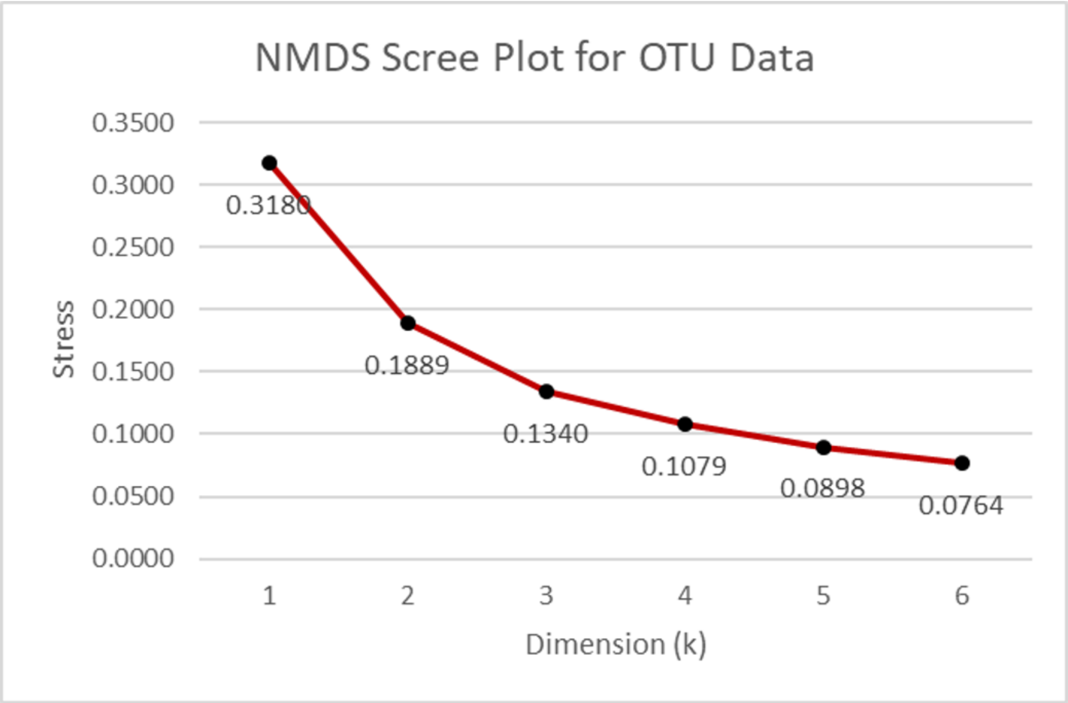


**Supplemental Figure 2.** NMDS scree plot of stress values for different levels of k calculated for a Bray-Curtis distance matrix of OTU community data that was transformed from read counts into percent abundance values prior to ordination.

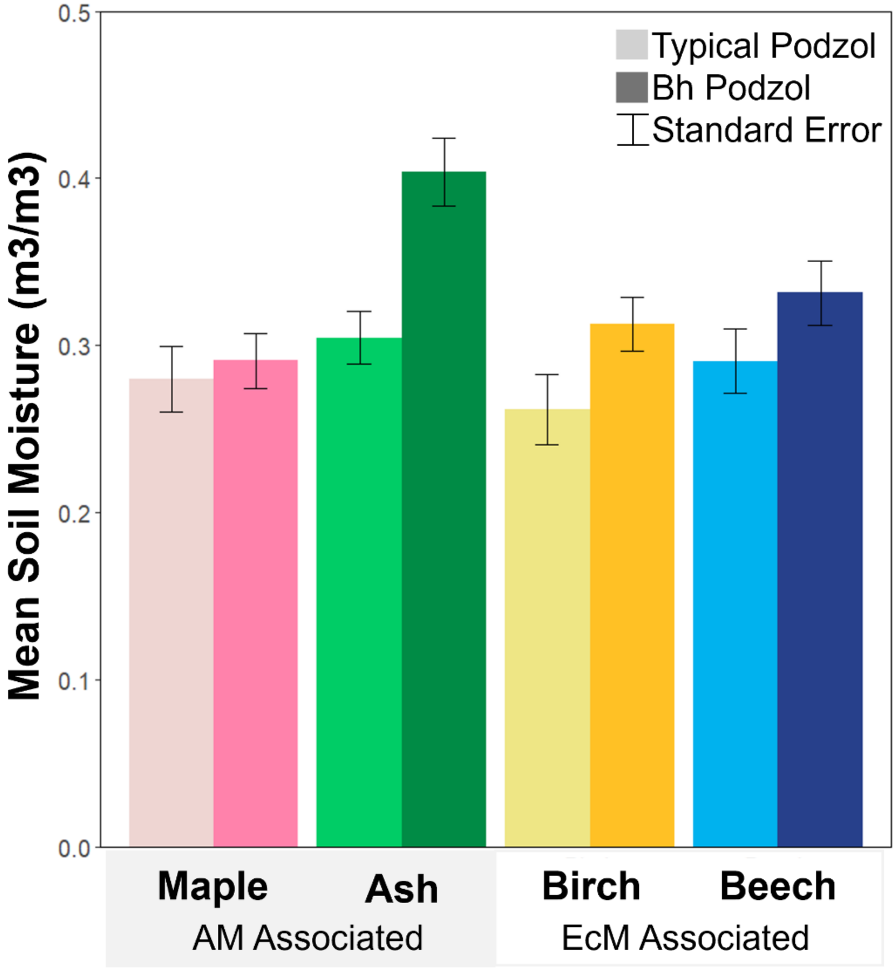


**Supplemental Figure 3.** Mean soil volumetric water content across four tree species and two soil types. Each combination of tree and soil type had 5 to 10 replicate plots and each of the 60 study plots had five moisture measurements recorded. Least square means were calculated by a three-way ANOVA which indicated differences in soil moisture across soil type and tree species with no significant cross interaction (Supplemental Table 1).

**Supplemental Tables**

**Supplemental Table 1.** Three-way ANOVA results comparing plot soil moisture across tree species, soil type, sampling occasion, and their cross interactions. Each of the 60 study plots had 5 measurements taken, which were combined in the ANOVA by treating plot labels as a random effect.

| Source | Degrees of Freedom | F statistic | p value |
|---|---|---|---|
| Soil Type | 1, 36 | 14.74 | 0.0005 |
| Tree Species | 3, 36 | 6.24 | 0.0016 |
| Sampling Occasion | 2, 36 | 12 | 0.0001 |
| Tree*Soil | 3, 36 | 2.04 | 0.1252 |
| Occasion*Soil | 2, 36 | 0.08 | 0.9223 |
| Occasion*Tree | 6, 36 | 0.81 | 0.5661 |
| Occasion*Tree*Soil | 6, 36 | 1.34 | 0.2657 |

**Supplemental Table 2A-1.** First half of the 90 Indicator OTUs for white ash (*Fraxinus americana*). OTU identity = this study's OTU ID number followed by the scientific name.

| Taxon Label | A Component | B Component | Indicator Value | p Value |
|---|---|---|---|---|
| OTU_1164_Gryganskiella_cystojenkinii | 0.9329 | 0.7333 | 0.827 | 0.001 |
| OTU_804_Unknown_Ascomycota | 0.8711 | 0.7333 | 0.799 | 0.001 |
| OTU_441_Hymenoscyphus_tamaricis | 0.8067 | 0.7333 | 0.769 | 0.001 |
| OTU_173_Unknown_Ascomycota | 0.9422 | 0.6 | 0.752 | 0.001 |
| OTU_234_Unknown_Ascomycota | 0.9054 | 0.6 | 0.737 | 0.002 |
| OTU_376_Mortierella_sp. | 0.9023 | 0.6 | 0.736 | 0.001 |
| OTU_481_Linnemannia_amoeboidea | 0.8731 | 0.6 | 0.724 | 0.002 |
| OTU_927_Alatospora_sp. | 0.9759 | 0.5333 | 0.721 | 0.001 |
| OTU_473_Saccharomycopsis_guyanensis | 0.6839 | 0.7333 | 0.708 | 0.002 |
| OTU_84_Tolypocladium_inflatum | 0.7024 | 0.6667 | 0.684 | 0.018 |
| OTU_1622_Unknown_Dothideomycetes | 0.8698 | 0.5333 | 0.681 | 0.001 |
| OTU_306_Unknown_Agaricomycetes | 0.9557 | 0.4667 | 0.668 | 0.015 |
| OTU_36_Unknown_Ascomycota | 0.477 | 0.9333 | 0.667 | 0.034 |
| OTU_509_Pleotrichocladium_opacum | 0.6062 | 0.7333 | 0.667 | 0.002 |
| OTU_1947_Anthopsis_sp. | 0.9069 | 0.4667 | 0.651 | 0.002 |
| OTU_865_Fontanospora_fusiramosa | 0.6915 | 0.6 | 0.644 | 0.011 |
| OTU_1010_Leucosporidium_sp. | 0.7711 | 0.5333 | 0.641 | 0.002 |
| OTU_2029_Unknown_Ascomycota | 0.8699 | 0.4667 | 0.637 | 0.001 |
| OTU_626_Acrophialophora_levis | 0.671 | 0.6 | 0.634 | 0.009 |
| OTU_123_Unknown_Ascomycota | 0.9857 | 0.4 | 0.628 | 0.018 |
| OTU_1078_Unknown_Ascomycota | 0.7332 | 0.5333 | 0.625 | 0.002 |
| OTU_328_Unknown_Ascomycota | 0.9713 | 0.4 | 0.623 | 0.007 |
| OTU_1529_Luellia_cystidiata | 0.9642 | 0.4 | 0.621 | 0.005 |
| OTU_585_Beauveria_caledonica | 0.819 | 0.4667 | 0.618 | 0.019 |
| OTU_719_Oidiodendron_sp. | 0.8174 | 0.4667 | 0.618 | 0.005 |
| OTU_202_Leucoglossum_leucosporum | 0.925 | 0.4 | 0.608 | 0.005 |
| OTU_1701_Unknown_Basidiomycota | 0.9225 | 0.4 | 0.607 | 0.001 |
| OTU_256_Penicillium_dodgei | 0.9183 | 0.4 | 0.606 | 0.003 |
| OTU_413_Unknown_Ascomycota | 0.5507 | 0.6667 | 0.606 | 0.007 |
| OTU_794_Pleuroascus_rectipilus | 0.7789 | 0.4667 | 0.603 | 0.006 |
| OTU_2652_Entoloma_luteofuscum | 0.9074 | 0.4 | 0.602 | 0.002 |
| OTU_881_Unknown_Basidiomycota | 0.7629 | 0.4667 | 0.597 | 0.006 |
| OTU_1048_Unknown_Hypocreales | 0.6598 | 0.5333 | 0.593 | 0.006 |
| OTU_1409_Unknown_Ascomycota | 0.8392 | 0.4 | 0.579 | 0.007 |
| OTU_1153_Unknown_Ascomycota | 1 | 0.3333 | 0.577 | 0.002 |
| OTU_3674_Ochrocladosporium_frigidarii | 1 | 0.3333 | 0.577 | 0.005 |
| OTU_5481_Alternaria_sp. | 1 | 0.3333 | 0.577 | 0.004 |
| OTU_3585_Unknown_Sordariomycetes | 0.9929 | 0.3333 | 0.575 | 0.003 |
| OTU_119_Unknown_Ascomycota | 0.8249 | 0.4 | 0.574 | 0.004 |
| OTU_1197_Unknown_Ascomycota | 0.6918 | 0.4667 | 0.568 | 0.006 |
| OTU_122_Camarophyllopsis_sp. | 0.9526 | 0.3333 | 0.564 | 0.023 |
| OTU_576_Unknown_Ascomycota | 0.7911 | 0.4 | 0.563 | 0.011 |
| OTU_1281_Entoloma_alboumbonatum | 0.7859 | 0.4 | 0.561 | 0.025 |
| OTU_158_Unknown_Ascomycota | 0.7824 | 0.4 | 0.559 | 0.022 |
| OTU_1434_Unknown_Ascomycota | 0.7733 | 0.4 | 0.556 | 0.01 |

**Supplemental Table 2A-2.** continued

| Taxon Label | A Component | B Component | Indicator Value | p Value |
|---|---|---|---|---|
| OTU_3689_Alatospora_sp. | 0.8913 | 0.3333 | 0.545 | 0.011 |
| OTU_1556_Unknown_Helotiales | 0.87 | 0.3333 | 0.539 | 0.015 |
| OTU_1273_Unknown_Ascomycota | 0.8629 | 0.3333 | 0.536 | 0.009 |
| OTU_4845_Exidia_glandulosa | 0.8574 | 0.3333 | 0.535 | 0.013 |
| OTU_656_Unknown_Agaricomycetes | 0.6036 | 0.4667 | 0.531 | 0.04 |
| OTU_1577_Fusarium_nurragi | 0.8389 | 0.3333 | 0.529 | 0.023 |
| OTU_1032_Unknown_Ascomycota | 0.5965 | 0.4667 | 0.528 | 0.015 |
| OTU_643_Unknown_Ascomycota | 0.8114 | 0.3333 | 0.52 | 0.037 |
| OTU_627_Unknown_Ascomycota | 1 | 0.2667 | 0.516 | 0.012 |
| OTU_1355_Unknown_Ascomycota | 1 | 0.2667 | 0.516 | 0.015 |
| OTU_1746_Pleotrichocladium_opacum | 1 | 0.2667 | 0.516 | 0.004 |
| OTU_2082_Trichoderma_polysporum | 1 | 0.2667 | 0.516 | 0.012 |
| OTU_2166_Unknown_Ascomycota | 1 | 0.2667 | 0.516 | 0.014 |
| OTU_2453_Unknown_Ascomycota | 1 | 0.2667 | 0.516 | 0.008 |
| OTU_2525_Tolypocladium_sp. | 1 | 0.2667 | 0.516 | 0.011 |
| OTU_2845_Gliocladium_sp. | 1 | 0.2667 | 0.516 | 0.009 |
| OTU_2990_Unknown_Ascomycota | 1 | 0.2667 | 0.516 | 0.014 |
| OTU_2991_Unknown_Dothideomycetes | 1 | 0.2667 | 0.516 | 0.015 |
| OTU_3029_Spirosphaera_sp. | 1 | 0.2667 | 0.516 | 0.012 |
| OTU_3623_Brachysporium_sp. | 1 | 0.2667 | 0.516 | 0.011 |
| OTU_1715_Unknown_Ascomycota | 0.7924 | 0.3333 | 0.514 | 0.027 |
| OTU_1152_Unknown_Ascomycota | 0.9845 | 0.2667 | 0.512 | 0.018 |
| OTU_2222_Unknown_Ascomycota | 0.98 | 0.2667 | 0.511 | 0.017 |
| OTU_1592_Coleophoma_sp. | 0.6502 | 0.4 | 0.51 | 0.023 |
| OTU_2893_Unknown_Sordariomycetes | 0.9572 | 0.2667 | 0.505 | 0.011 |
| OTU_2295_Unknown_Sordariomycetes | 0.9545 | 0.2667 | 0.505 | 0.011 |
| OTU_934_Chaetosphaeria_hebetiseta | 0.9532 | 0.2667 | 0.504 | 0.042 |
| OTU_3164_Tuber_sp. | 0.9368 | 0.2667 | 0.5 | 0.019 |
| OTU_2109_Unknown_Ascomycota | 0.936 | 0.2667 | 0.5 | 0.024 |
| OTU_109_Unknown_Ascomycota | 0.9279 | 0.2667 | 0.497 | 0.046 |
| OTU_851_Unknown_Microbotryomycetes | 0.6056 | 0.4 | 0.492 | 0.036 |
| OTU_355_Cenococcum_sp. | 0.599 | 0.4 | 0.489 | 0.05 |
| OTU_2392_Unknown_Sordariomycetes | 0.8963 | 0.2667 | 0.489 | 0.016 |
| OTU_2685_Metarhizium_robertsii | 0.7027 | 0.3333 | 0.484 | 0.034 |
| OTU_2314_Unknown_Ascomycota | 0.8771 | 0.2667 | 0.484 | 0.019 |
| OTU_2749_Unknown_Ascomycota | 0.8685 | 0.2667 | 0.481 | 0.029 |
| OTU_3208_Unknown_Ascomycota | 0.8524 | 0.2667 | 0.477 | 0.027 |
| OTU_1707_Unknown_Ascomycota | 0.6736 | 0.3333 | 0.474 | 0.036 |
| OTU_979_Unknown_Ascomycota | 0.8299 | 0.2667 | 0.47 | 0.023 |
| OTU_1211_Gryganskiella_cystojenkinii | 0.6636 | 0.3333 | 0.47 | 0.042 |
| OTU_2102_Unknown_Ascomycota | 0.8278 | 0.2667 | 0.47 | 0.037 |
| OTU_4606_Unknown_Tremellodendropsidales | 1 | 0.2 | 0.447 | 0.044 |
| OTU_2778_Unknown_Leotiomycetes | 0.7335 | 0.2667 | 0.442 | 0.031 |
| OTU_5490_Unknown_Ascomycota | 0.9693 | 0.2 | 0.44 | 0.041 |
| OTU_314_Unknown_Ascomycota | 0.9604 | 0.2 | 0.438 | 0.047 |

**Supplemental Table 2B.** Indicator OTUs for yellow birch (*Betula alleghaniensis*).

OTU identity = this study's OTU ID number followed by the scientific name.

| Taxon Label | A Component | B Component | Indicator Value | p Value |
|---|---|---|---|---|
| OTU_329_Unknown_Ascomycota | 0.7418 | 1 | 0.861 | 0.001 |
| OTU_619_Didymella_sp. | 0.8198 | 0.8 | 0.81 | 0.002 |
| OTU_150_Penicillium_longicatenatum | 0.6599 | 0.9333 | 0.785 | 0.001 |
| OTU_702_Unknown_Agaricomycetes | 0.7273 | 0.8 | 0.763 | 0.001 |
| OTU_832_Unknown_Stachybotryaceae | 0.5817 | 0.8667 | 0.71 | 0.002 |
| OTU_307_Tolypocladium_sp. | 0.5328 | 0.9333 | 0.705 | 0.016 |
| OTU_181_Oidiodendron_sp. | 0.5693 | 0.8667 | 0.702 | 0.002 |
| OTU_494_Unknown_Pezizaceae | 0.7138 | 0.6667 | 0.69 | 0.002 |
| OTU_461_Unknown_Helotiales | 0.7015 | 0.6667 | 0.684 | 0.004 |
| OTU_485_Unknown_Leotiomycetes | 0.8566 | 0.5333 | 0.676 | 0.006 |
| OTU_261_Unknown_Hypocreales | 0.4522 | 1 | 0.672 | 0.017 |
| OTU_550_Unknown_Venturiaceae | 0.9589 | 0.4667 | 0.669 | 0.001 |
| OTU_278_Unknown_Sordariomycetes | 0.9234 | 0.4667 | 0.656 | 0.005 |
| OTU_2218_Unknown_Helotiales | 0.8035 | 0.5333 | 0.655 | 0.009 |
| OTU_1038_Setomelanomma_holmii | 0.8685 | 0.4667 | 0.637 | 0.014 |
| OTU_429_Oidiodendron_maius | 0.448 | 0.8667 | 0.623 | 0.009 |
| OTU_3212_Parapyrenochaeta_sp. | 0.9666 | 0.4 | 0.622 | 0.001 |
| OTU_2_Saitozyma_sp. | 0.3852 | 1 | 0.621 | 0.017 |
| OTU_46_Trichoderma_sp. | 0.3758 | 1 | 0.613 | 0.023 |
| OTU_488_Unknown_Helotiales | 0.5364 | 0.6667 | 0.598 | 0.018 |
| OTU_1544_Unknown_Ascomycota | 0.6492 | 0.5333 | 0.588 | 0.037 |
| OTU_299_Unknown_Cordycipitaceae | 0.4526 | 0.7333 | 0.576 | 0.041 |
| OTU_666_Penicillium_dodgei | 0.6625 | 0.4667 | 0.556 | 0.021 |
| OTU_1133_Cladophialophora_sp. | 0.5611 | 0.5333 | 0.547 | 0.023 |
| OTU_2013_Unknown_Ascomycota | 0.7414 | 0.4 | 0.545 | 0.025 |
| OTU_1993_Infundichalara_sp. | 0.8623 | 0.3333 | 0.536 | 0.022 |
| OTU_1546_Unknown_Ascomycota | 0.5713 | 0.4667 | 0.516 | 0.043 |
| OTU_822_Unknown_Ascomycota | 0.9878 | 0.2667 | 0.513 | 0.031 |
| OTU_446_Russula_redolens | 0.9859 | 0.2667 | 0.513 | 0.044 |
| OTU_3471_Unknown_Helotiaceae | 0.7834 | 0.3333 | 0.511 | 0.05 |
| OTU_2140_Unknown_Basidiomycota | 0.955 | 0.2667 | 0.505 | 0.015 |
| OTU_551_Unknown_Thelephoraceae | 0.9544 | 0.2667 | 0.504 | 0.029 |
| OTU_2327_Sympodiella_sp. | 0.9469 | 0.2667 | 0.503 | 0.033 |
| OTU_1494_Unknown_Hypocreales | 0.6243 | 0.4 | 0.5 | 0.019 |
| OTU_3262_Phaeotremella_fagi | 0.9305 | 0.2667 | 0.498 | 0.043 |
| OTU_1927_Polyozellus_tristis | 1 | 0.2 | 0.447 | 0.04 |
| OTU_4753_Basidiodendron_sp. | 1 | 0.2 | 0.447 | 0.047 |

**Supplemental Table 2C.** Indicator OTUs for American beech (*Fagus grandifolia*).

OTU identity = this study's OTU ID number followed by the scientific name.

| Taxon Label | A Component | B Component | Indicator Value | p Value |
|---|---|---|---|---|
| OTU_617_Unknown_Basidiomycota | 0.861 | 0.7333 | 0.795 | 0.001 |
| OTU_1362_Haptocillium_sp. | 0.9354 | 0.5333 | 0.706 | 0.001 |
| OTU_1755_Unknown_Sordariomycetes | 0.8223 | 0.6 | 0.702 | 0.001 |
| OTU_251_Unknown_Chaetosphaeriaceae | 0.581 | 0.8 | 0.682 | 0.009 |
| OTU_1165_Unknown_Ascomycota | 0.7571 | 0.6 | 0.674 | 0.002 |
| OTU_80_Mortierella_pulchella | 0.4745 | 0.9333 | 0.666 | 0.013 |
| OTU_533_Unknown_Hypocreales | 0.803 | 0.5333 | 0.654 | 0.035 |
| OTU_209_Unknown_Basidiomycota | 0.6715 | 0.6 | 0.635 | 0.022 |
| OTU_1264_Mortierella_turficola | 0.6513 | 0.6 | 0.625 | 0.001 |
| OTU_597_Unknown_Dermateaceae | 0.35 | 1 | 0.592 | 0.041 |
| OTU_1391_Unknown_Helotiales | 0.6454 | 0.5333 | 0.587 | 0.012 |
| OTU_1059_Unknown_Hypocreales | 0.7358 | 0.4667 | 0.586 | 0.04 |
| OTU_349_Unknown_Helotiales | 0.7345 | 0.4667 | 0.585 | 0.018 |
| OTU_1971_Unknown_Basidiomycota | 0.6271 | 0.5333 | 0.578 | 0.01 |
| OTU_768_Unknown_Sordariales | 0.9971 | 0.3333 | 0.577 | 0.033 |
| OTU_715_Unknown_Sordariomycetes | 0.9895 | 0.3333 | 0.574 | 0.005 |
| OTU_1973_Unknown_Ascomycota | 0.6818 | 0.4667 | 0.564 | 0.012 |
| OTU_3002_Hirsutella_rhossiliensis | 0.8847 | 0.3333 | 0.543 | 0.007 |
| OTU_397_Oidiodendron_maius | 0.5418 | 0.5333 | 0.538 | 0.044 |
| OTU_1901_Unknown_Endogonomycetes_GS21 | 0.6111 | 0.4667 | 0.534 | 0.029 |
| OTU_1966_Unknown_Basidiomycota | 0.7064 | 0.4 | 0.532 | 0.014 |
| OTU_1367_Unknown_Sordariomycetes | 0.6774 | 0.4 | 0.521 | 0.049 |
| OTU_1042_Unknown_Mortierellaceae | 0.6741 | 0.4 | 0.519 | 0.013 |
| OTU_408_Trichoderma_alni | 0.5743 | 0.4667 | 0.518 | 0.035 |
| OTU_1943_Unknown_Endogonomycetes_GS21 | 0.6293 | 0.4 | 0.502 | 0.032 |
| OTU_1689_Trichoderma_sp. | 0.93 | 0.2667 | 0.498 | 0.032 |
| OTU_280_Russula_emetica | 0.9008 | 0.2667 | 0.49 | 0.029 |
| OTU_1618_Unknown_Basidiomycota | 0.8979 | 0.2667 | 0.489 | 0.049 |
| OTU_1191_Unknown_Rozellomycota | 0.8722 | 0.2667 | 0.482 | 0.05 |
| OTU_2274_Unknown_Agaricomycetes | 1 | 0.2 | 0.447 | 0.038 |
| OTU_2850_Unknown_Basidiomycota | 1 | 0.2 | 0.447 | 0.05 |
| OTU_1429_Unknown_Stachybotryaceae | 0.98 | 0.2 | 0.443 | 0.05 |
| OTU_1608_Cortinarius_boulderensis | 0.8619 | 0.2 | 0.415 | 0.048 |

**Supplemental Table 2D.** Indicator OTUs for sugar maple (*Acer saccharum*).

OTU identity = this study's OTU ID number followed by the scientific name.

| Taxon Label | A Component | B Component | Indicator Value | p Value |
|---|---|---|---|---|
| OTU_628_Sympodiella_acicola | 0.5367 | 0.9333 | 0.708 | 0.001 |
| OTU_755_Unknown_Basidiomycota | 0.5027 | 0.8667 | 0.66 | 0.02 |
| OTU_167_Unknown_Ascomycota | 0.5608 | 0.7333 | 0.641 | 0.043 |
| OTU_71_Ramariopsis_sp. | 0.6785 | 0.6 | 0.638 | 0.047 |
| OTU_753_Unknown_Helotiales | 0.7393 | 0.5333 | 0.628 | 0.006 |
| OTU_1316_Pilidium_sp. | 0.8436 | 0.4667 | 0.627 | 0.022 |
| OTU_99_Hygrocybe_flavescens | 0.9946 | 0.3333 | 0.576 | 0.015 |
| OTU_1909_Unknown_Helotiales | 0.7904 | 0.4 | 0.562 | 0.009 |
| OTU_680_Unknown_Venturiales | 0.9441 | 0.3333 | 0.561 | 0.032 |
| OTU_1398_Unknown_Ascomycota | 0.6485 | 0.4667 | 0.55 | 0.021 |
| OTU_1827_Unknown_Leotiomycetes | 0.5577 | 0.5333 | 0.545 | 0.025 |
| OTU_1621_Lecanicillium_fusisporum | 0.8735 | 0.3333 | 0.54 | 0.011 |
| OTU_989_Collarina_sp. | 0.8713 | 0.3333 | 0.539 | 0.032 |
| OTU_1007_Chaetosphaeria_myriocarpa | 0.8099 | 0.3333 | 0.52 | 0.02 |
| OTU_1917_Entoloma_luteofuscum | 0.6697 | 0.4 | 0.518 | 0.029 |
| OTU_3298_Unknown_Sordariomycetes | 1 | 0.2667 | 0.516 | 0.013 |
| OTU_3475_Unknown_Leotiomycetes | 1 | 0.2667 | 0.516 | 0.012 |
| OTU_939_Unknown_Basidiomycota | 0.9974 | 0.2667 | 0.516 | 0.014 |
| OTU_402_Cortinarius_sp. | 0.9905 | 0.2667 | 0.514 | 0.018 |
| OTU_2064_Unknown_Basidiomycota | 0.6126 | 0.4 | 0.495 | 0.044 |
| OTU_1969_Unknown_Ascomycota | 0.7261 | 0.3333 | 0.492 | 0.044 |
| OTU_1590_Unknown_Eurotiomycetes | 0.6682 | 0.3333 | 0.472 | 0.045 |
| OTU_2719_Unknown_Mycosphaerellales | 1 | 0.2 | 0.447 | 0.047 |
| OTU_5373_Unknown_Micropeltidaceae | 1 | 0.2 | 0.447 | 0.048 |
| OTU_4451_Unknown_Phacidiaceae | 0.9596 | 0.2 | 0.438 | 0.05 |
| OTU_4005_Unknown_Sordariomycetes | 0.712 | 0.2667 | 0.436 | 0.046 |
